# A spatial map of human liver cirrhosis reveals the patho-architecture and gene signatures associated with cell state transitions during liver disease

**DOI:** 10.1101/2023.06.28.546905

**Authors:** Nigel L Hammond, Syed Murtuza Baker, Sokratia Georgaka, Ali Al-Anbaki, Elliot Jokl, Kara Simpson, Varinder S Athwal, Ajith K Siriwardena, Harry VM Spiers, Mike J Dixon, Nicoletta Bobola, Andrew D Sharrocks, Neil A Hanley, Magnus Rattray, Karen Piper Hanley

## Abstract

Liver fibrosis is a major cause of death worldwide. As a progressive step in chronic liver disease, fibrosis is almost always diagnosed too late with limited treatment options. Here, we uncover the spatial transcriptional landscape driving human liver fibrosis using single nuclei RNA and Assay for Transposase-Accessible Chromatin (ATAC) sequencing to deconvolute multi-cell spatial transcriptomic profiling in human liver cirrhosis. Through multi-modal data integration, we define molecular signatures driving cell state transitions in liver disease and define an impaired cellular response and directional trajectory from hepatocytes to cholangiocytes associated with disease remodelling. We identify pro-fibrogenic signatures in non-parenchymal cell subpopulations co-localised within the fibrotic niche and localise transitional cell states at the scar interface. This combined approach provides a spatial atlas of gene regulation and defines molecular signatures associated liver disease for targeted therapeutics or as early diagnostic markers of progressive liver disease.

## Introduction

Chronic liver disease is increasing and a major cause of death worldwide^1^. Although the liver has a remarkable capacity to regenerate after acute and transient injuries, repeated insult leads to progressive scarring (or fibrosis) and ultimately end-stage cirrhosis requiring transplant^2, 3^. Despite major advances in recent years, diagnosis tends to occur in advanced disease and clinically approved antifibrotic treatments are severely limited. A deeper understanding of the spatial complexity driving molecular and cellular mechanisms in human liver disease will be critical to identify early changes in pathophysiology leading to progressive disease and to develop novel therapeutics.

Pathological fibrosis underlies all forms chronic liver disease and is associated with major changes to both the quantity and composition of the extracellular matrix (ECM)^4^. After acute liver injury, parenchymal cells (predominantly hepatocytes) regenerate and stimulate an inflammatory response^3^. Subsequently, hepatic stellate cells (HSCs), the main ECM-producing cells of the injured liver, become activated and transdifferentiate into pro-fibrotic myofibroblasts^4^. Iterative injury prolongs an impaired wound-healing response, resulting in the accumulation of excess ECM and the formation of fibrous scar tissue. Over time profound changes to the liver parenchyma and vasculature progress to cirrhosis, characterised by the formation of fibrous septa and regenerative nodules^4^. This unique microenvironment, involving complex interactions between multiple cell-types within the fibrotic niche, creates an impaired regenerative response and progressive disease^2, 3, 4^.

To advance understanding into the heterogeneity of liver, single cell and single nuclei RNA sequencing (sc/snRNA-seq) have uncovered gene signatures associated within cell populations to define progenitor cells, gene expression associated with zonation, and cellular response to disease pathogenesis^5, 6, 7, 8, 9^. Although exciting, dissociated tissue involved in scRNA-seq can cause transcriptomic changes^10^ and crucial information on the spatial context of cells is lost. In contrast, spatially resolved transcriptomic (ST) technologies^11^ have enabled genome-wide characterisation of cellular heterogeneity at near single-cell resolution, while preserving spatial information. In human liver, ST studies have defined structural zonation in normal liver and hepatic macrophage states in disease^12, 13, 14, 15^. While all are important developments to the field, the impaired regenerative response of parenchymal cells and role of the local scar environment altering cell state has been overlooked.

Here, we integrate ST, snRNA-seq and chromatin accessibility to characterise the spatial transcriptional landscape in human cirrhotic liver. We uncover the cellular patho-architecture of diseased liver and define spatial molecular signatures of fibrotic scars, damaged parenchyma and impaired regenerative response at the scar interface. Through pseudo-temporal ordering, we show cell state transitions and directional trajectory from hepatocytes to cholangiocytes, potentially demarcating impaired regeneration. We infer the gene regulatory networks (GRNs) differentiating these cell states to gain novel insight into the gene signatures and regulatory programmes driving an impaired regenerative response during liver disease. Together, these data serve as a spatial atlas of gene regulation and altered cell state in human cirrhosis and provides a valuable resource to explore biomarkers and potential therapeutic targets in progressive liver disease.

## Results

### ST resolves the spatial signature of fibrotic scars and altered cell state at their interface inhuman liver cirrhosis

Fresh, unfixed human cirrhotic tissue from three donors were collected to establish spatially resolved datasets using the 10X Visium platform (Supplementary Information, Supplementary Figures 1A and 2, Supplementary Table 1). Following processing and mapping, histological landmarks were apparent demarcating fibrous scar from the surrounding liver parenchyma. As expected for human samples the degree of fibrosis varied, ranging from numerous fibrous septa (sample a) to extensive bridging septa encompassing regenerative nodules (sample b) and localised portal fibrosis (sample c) (Supplementary Figure 1B-D).

To explore gene expression underlying the cirrhotic samples, dimensionality reduction of spatial data was performed followed by unsupervised clustering and differential gene analysis (Supplementary Information and Supplementary Figure 1). However, to capture the most diverse cell-types and provide broad insight into spatial gene expression associated with cirrhosis we further interrogated sample b, displaying the most diversity in cell type clusters (Supplementary Figure 1), and compared analysis using both *k-*means and unsupervised Louvain clustering methods (Figure 1A-D, Supplementary Tables 2 and 3). While *k*-means defined the spatial heterogeneity of fibrotic scars (Figure 1A, B and Supplementary Table 2), Louvain clustering defined heterogeneity within the functioning liver parenchyma (b7, b8, b10) (Figure 1C, D and Supplementary Table 3). In particular, cluster b9 (red) defined the tissue-interface with the fibrotic scar (b6; orange) potentially associated with damaged or impaired / regenerating parenchyma (Figure 1C, D). For both analyses, we displayed differentially expressed genes (DEGs) per cluster as a heatmap, and annotated co-expression gene modules to spatial regions within the tissue (e.g. scar, interface, tissue; Figure 1B, D).

**Figure 1.**
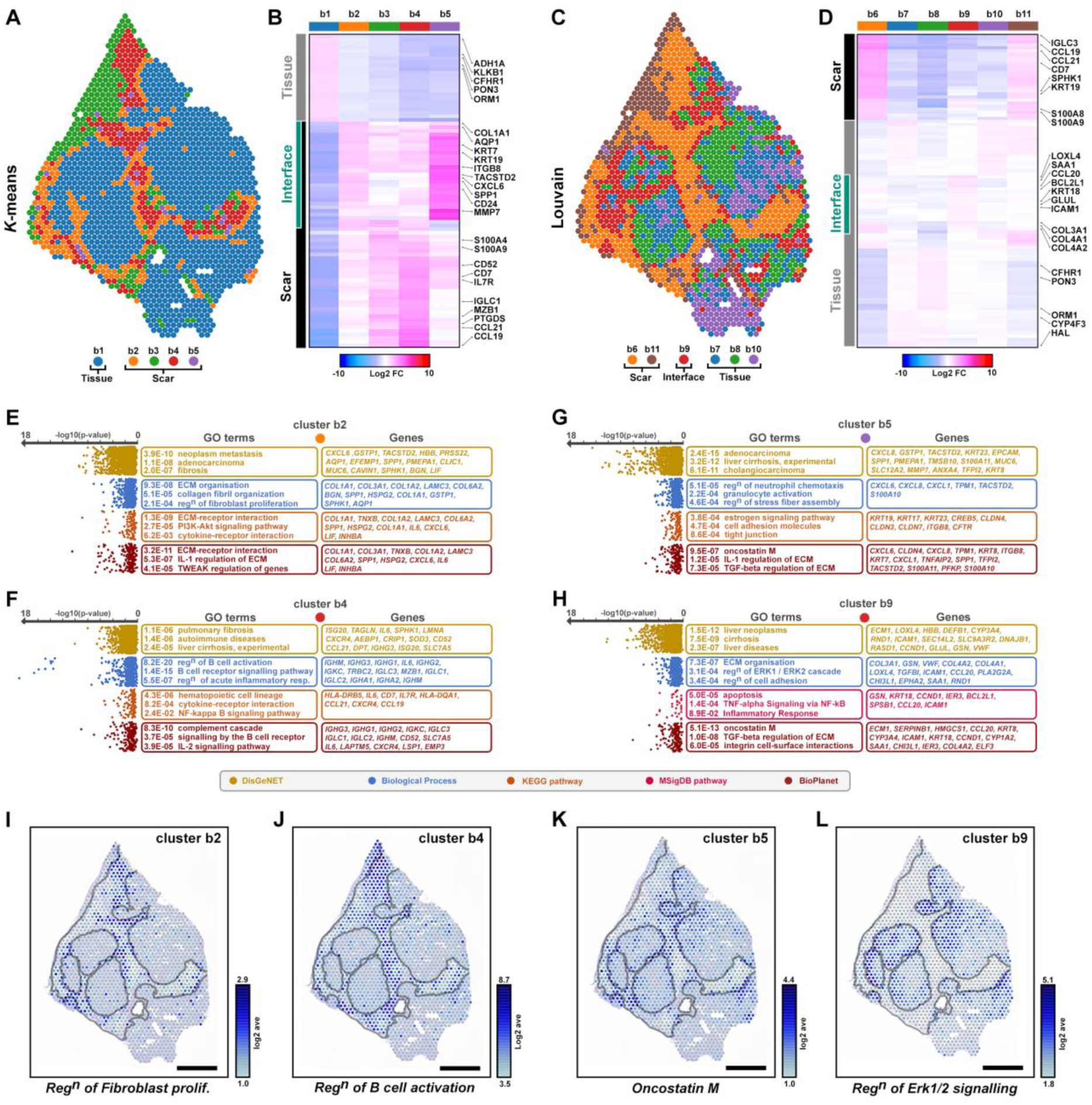
Spatial signatures underlying fibrotic tissue and interface with liver parenchyma. (**A**) Visualisation of spatial transcriptomic clusters for k-means (**A**) shows heterogeneity within fibrotic scar, identifies scar and interface clusters, with significant differentially expressed genes shown as a heatmap (**B**). Louvain clustering shows heterogeneity within liver parenchyma (**C**), including a tissue-interface cluster and significant genes underlying them (**D**). (**E**) Gene ontology (GO) of significant genes underlying clusters. (**F-I**) Spatial expression of gene modules underlying GO terms with corresponding spatial expression of select genes. Scale bars, 1mm.

Gene ontology (GO; Supplementary Table 4) analysis of *k*-means clusters identified *fibrosis*, *regulation of fibroblast proliferation* and *ECM-receptor interaction* terms within the scar interface cluster (b2; Figure 1E) that spatially mapped to the periphery of the scar (Figure 1I). As expected, pro-fibrotic markers and ECM components associated with activated HSCs were also evident and enriched in cluster b2 (*COL1A1*, *COL1A2*, *COL3A1*, *COL6A2*, *TIMP1*, *BGN*, *LAMC3*, *ADAMTSL2*, *SPHK1*, *AEBP1*, *CCN2*), with several targets co-expressed at the scar interface (Supplementary Figure 4A). Scar-associated cluster b4 was significantly enriched for *B cell signalling* and *B cell activation*, mapping spatially to the centre of the scar (Figure 1F,J). Accordingly, marker genes of immune and activated B cells (*IGLC2*, *IGHG1*, *IGHG3*, *JCHAIN, MZB1*, *DERL3, S100A9*, *LSP1*) were enriched in this region, along with several mechano-sensitive components (*LMNA*, *TALGN*) (Figure 1F; Supplementary Figure 4B). Although cluster b5 was the smallest cluster (28/1609 spots), these spots occupied discrete locations at the scar interface. GO analysis revealed enrichment for genes involved in *cholangiocarcinoma*, *tight junction assembly* and *oncostatin m* signalling, which also spatially mapped to discrete regions within and towards the edge of the scar (Figure 1G, K). Markers associated with cholangiocytes (*KRT19*, *KRT7*, *KRT23*, *EPCAM*, *AQP1*, *SPP1, CD24*) and chemokine ligands (*CXCL1*, *CXCL6*, *CXCL8*), were enriched both in cluster b2 and b5, localised at the scar interface (Figure 1E,G,K; Supplementary Figure 4C).

Similarly, GO analysis of DEGs from Louvain clustering indicated enrichment of genes within cluster b9 (red; Figure XX) were involved in *ERK1/2 signalling*, *apoptosis*, *inflammatory response* and *integrin cell surface interactions* (Figure 1H) spatially located at the scar interface (Figure 1L). These regions also correlated with the spatial expression of pro-fibrotic ECM components (*LOXL4*, *COL3A1*, *COL4A1*), damage-induced inflammatory response proteins (*SAA1*, *SAA2*, *DEF1B*), apoptosis (*GDF*-*15*, *BCL2L1*, *TM4SF1*), and chemokine markers associated with hepatocyte regeneration (*CCL20, ICAM-1*) (Figure 1H; Supplementary Figure 4D).

Taken together, ST uncovered gene signatures underlying the pathophysiology in human cirrhotic liver tissue and altered cell states demarcating an impaired response to on-going disease. However, information on the cell-type contribution to these signatures was obfuscated by the current resolution of the spatial spots.

### Integrated spatial multi-omic analyses of human cirrhotic liver

To improve resolution and provide a refined spatial cell-type map, we jointly profiled the transcriptome (snRNA-seq) and accessible chromatin (snATAC-seq) from human cirrhotic liver at single nuclei resolution (from sample b Figure 2). From 6,170 nuclei, an average of 2,445 genes and 8,368 UMIs per nuclei were detected for snRNA-seq (Supplementary Figure 5A, B). Following dimensionality reduction^16^ and low resolution clustering, seven hepatic cell populations were identified (Figure 2B, C). In combination with publicly available datasets^5, 7, 9, 17^, differential expression analyses identified markers for all major liver cell-types in our dataset, specifically hepatocytes (HC; 45%), endothelial cells (EC; 26%), cholangiocytes (CC; 9%), mesenchymal cells (MC; 7%), macrophages (MP; 6%), lymphocytes (LC; 6.5%) and B cell (BC; 0.5%) populations (Figure 2B, C). From the entire dataset we further uncovered cellular heterogeneity to define 19 subclusters within the seven discrete cell populations (Figure 2B, C; and Supplementary Table 5).

**Figure 2.**
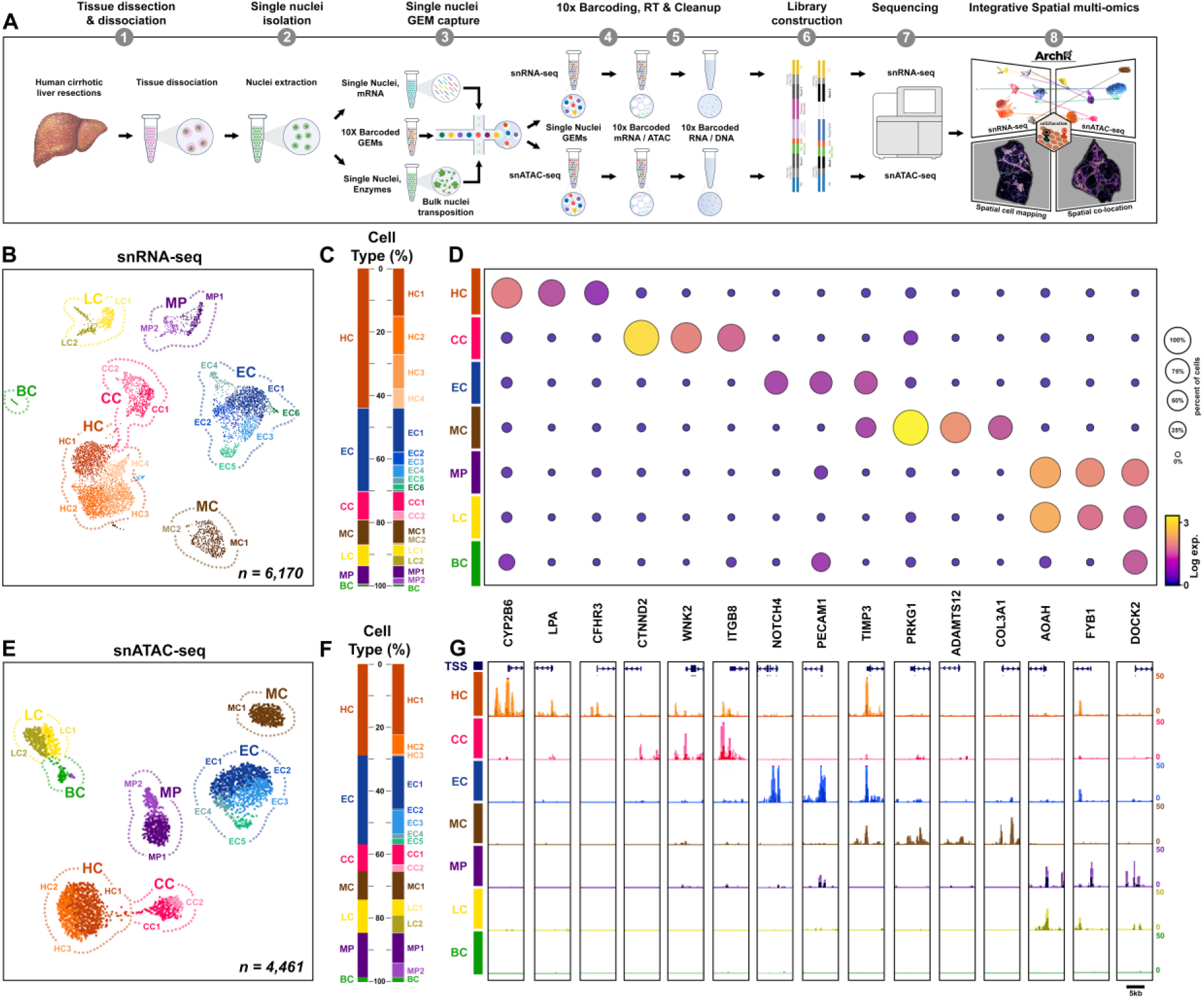
Integrated single nuclei multi-omic analysis of human cirrhotic liver. (**A**) Schematic of 10X Chromium workflow to generate snRNA-seq and snATAC-seq libraries from cirrhotic liver resections, followed by sequencing and integrated spatial multi-omic analysis. (**B**) UMAP clusters shows 7 discrete cell populations, which when re-clustered at high resolution reveal 19 sub-clusters within lineages. (**C**) Stacked bar chart shows the proportion of nuclei assigned to each cluster/sub-cluster. (**D**) Dot plot shows transcript expression and cell proportions of marker genes across cell types. (**E**) UMAP plot of snATAC-seq cell types with label transfer of sub-clusters from snRNA-seq data and (**F**) proportion of nuclei within each cluster/subcluster. (**G**) Promoter accessibility of cell type marker genes and correlation with gene expression (**D**). HC, hepatocytes; CC, cholangiocytes; EC, endothelial cells; MC, mesenchymal cells; MP, macrophages; LC, lymphocytes; BC, b cells.

Simultaneously, we profiled chromatin accessibility from the same human cirrhotic liver sample. From 4,461 nuclei, we detected 199,455 regions of accessible chromatin which using unsupervised clustering resolved to seven discrete cell populations (Figure 2E, F and and Supplementary Table 6). To define cell-types based on chromatin accessibility profiles, we leveraged our annotated snRNA-seq data and performed label transfer using Seurat^18^. Integration of paired modalities confirmed that all major liver cell-types were present in both datasets (Figure 2C, F). Furthermore, transfer of snRNA-seq subcluster annotations confirmed the majority of cell states (16 out of 19 subclusters) were represented in the snATAC-seq dataset (Figure 2B, C, E, F). Combined analysis of cell-type markers of gene expression and promotor accessibility across both modalities were highly correlated, validating our integrative single nuclei multi-omic approach (Figure 2D, G and Supplementary Tables 5-6).

### Spatial heterogeneity of non-parenchymal cell populations in human cirrhotic liver

Next, we explored snRNA-seq gene signatures underlying the heterogeneity of non parenchymal cell-types in cirrhotic liver. ECs were identified by the marker genes *PTPRB*, *LDB2*, *NRG3*, *NOSTRIN* and *AKAP12*, and were further characterised into six distinct sub-clusters (EC1-6) (Figure 3A). EC1, 2 and 3 represented over 80% of the total ECs and were annotated as liver sinusoidal endothelial cells (LSECs; a specialised microvascular EC) using the marker genes (*RELN*, *STAB1*, *STAB2*, *MRC1*, *CD36*) (Figure 3B, C). Although clusters EC4 and EC5 were well separated in UMAP, they uniquely co-expressed several genes (*ANO2*, *VWF*, *CARMIL1*, *BMX*, *CDH23*) indicating a vascular EC phenotype. Specifically, EC4 was enriched for WNT components and other markers (*RSPO3*, *WNT9B*, *WNT2*, *LHX6*, *CELF4*) suggesting this population were central venous ECs, while EC5 marker genes (*SNCAIP*, *CDH11*, *MYOF*, *NAV3*, *TNFRSF11A*) identified these cells as portal ECs (Figure 3A-C). Cluster EC6 consisted of just 31 cells and expressed markers of proliferating LSECs (*MKI67*, *PCNA, RELN*). Spatial mapping of EC subpopulations revealed the distribution of LSECs (EC1-3) within the main tissue architecture, while EC4-6 were differentially enriched within regions of the scar, suggesting a shared profibrotic signature (Figure 3D). Of interest, we have previously described NAV3 (from EC5) in activated pericytes (kidney specific myofibroblasts) during chronic kidney disease^19^ highlighting potentially shared mechanisms of fibrosis.

**Figure 3.**
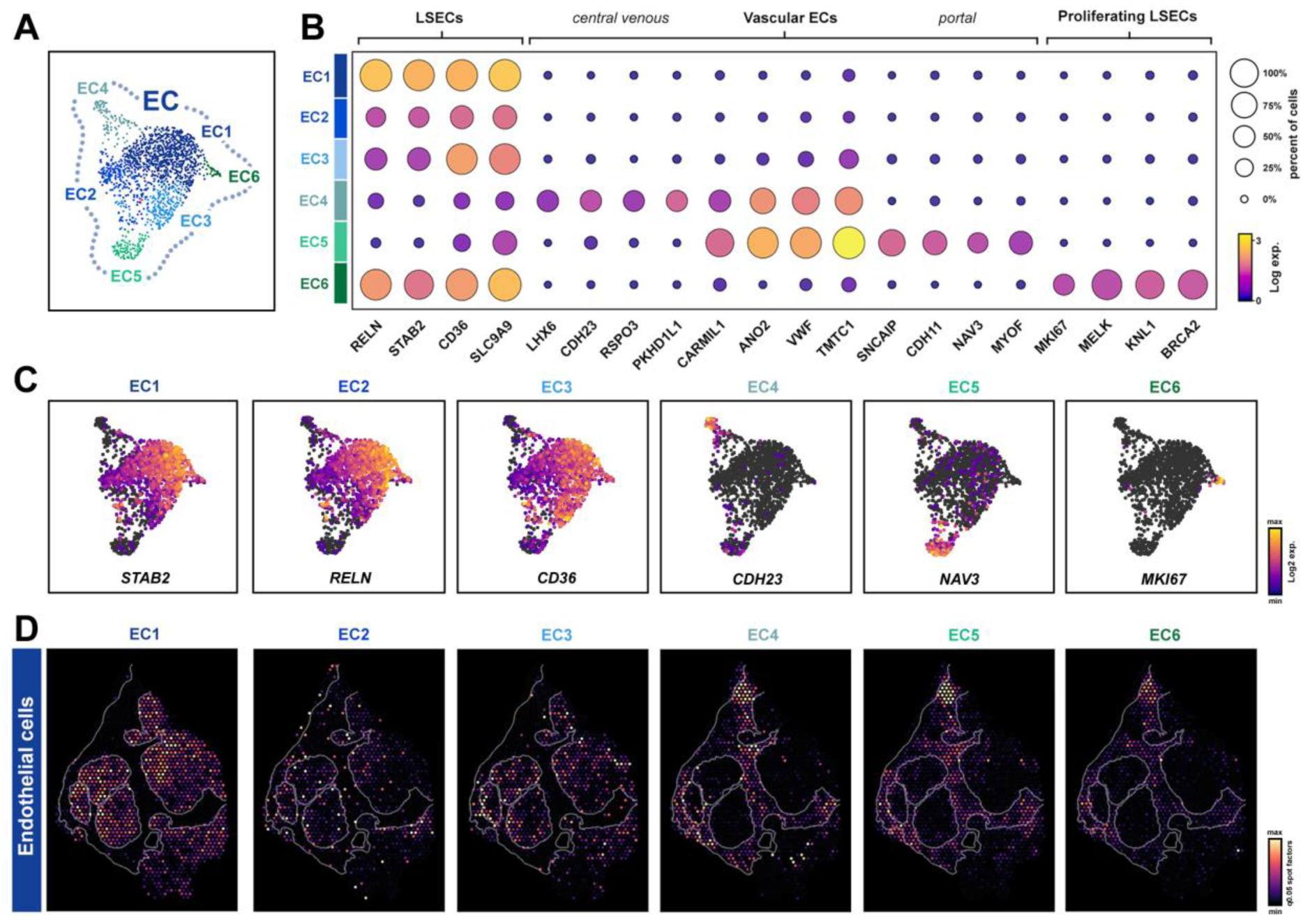
Spatial heterogeneity of endothelial cell populations in cirrhotic liver. (**A**-**C**) UMAP sub-clusters, dot plots and marker gene expression plots showing heterogeneity of sub-populations of endothelial cells (EC). (**D**) Spatial deconvolution maps of EC sub-populations within cirrhotic sample b. LSECs, liver sinusoidal endothelial cells.

MCs displayed markers representative of resident liver fibroblasts, Hepatic Stellate Cells (HSCs), which in response to liver injury can transdifferentiate into profibrotic activated myofibroblasts. MC1 comprised the majority of MCs (Figure 4A) and was enriched for *PDGFRA* and *PDGFRB*, alongside HSC marker genes (*LAMA2*, *ADAMTS12*, *ADAMTSL1*, *RELN*, *VIPR1, NAV3*) (Figure 4A, B, G). Consistent with previous studies^6, 9, 20^, MC1 displayed markers of quiescent qHSCs present in healthy liver (*RBP1*, *LRAT*, *SPARCL1*, *LHX2*) and activated HSCs (aHSCs) responsible for tissue damaging ECM deposition (*COL1A1*, *TIMP1*, *ACTA2*, *NCAM*, *SPARC*). MC2 comprised only 17 cells and were *PDGFRA*-/*PDGFRB*+, indicating a vascular smooth muscle cells (vSMCs) origin. As expected, differential gene analysis revealed marker genes of vSMCs (*MYH11*, *MCAM*, *NOTCH3*, *PPARG, CNN1*) enriched in MC2 (Figure 4A, B, G). Spatially, MC1 and MC2 mapped to discrete regions within the scar (Figure 4J) with a similar profile to EC4-6 (Figure 3D).

**Figure 4.**
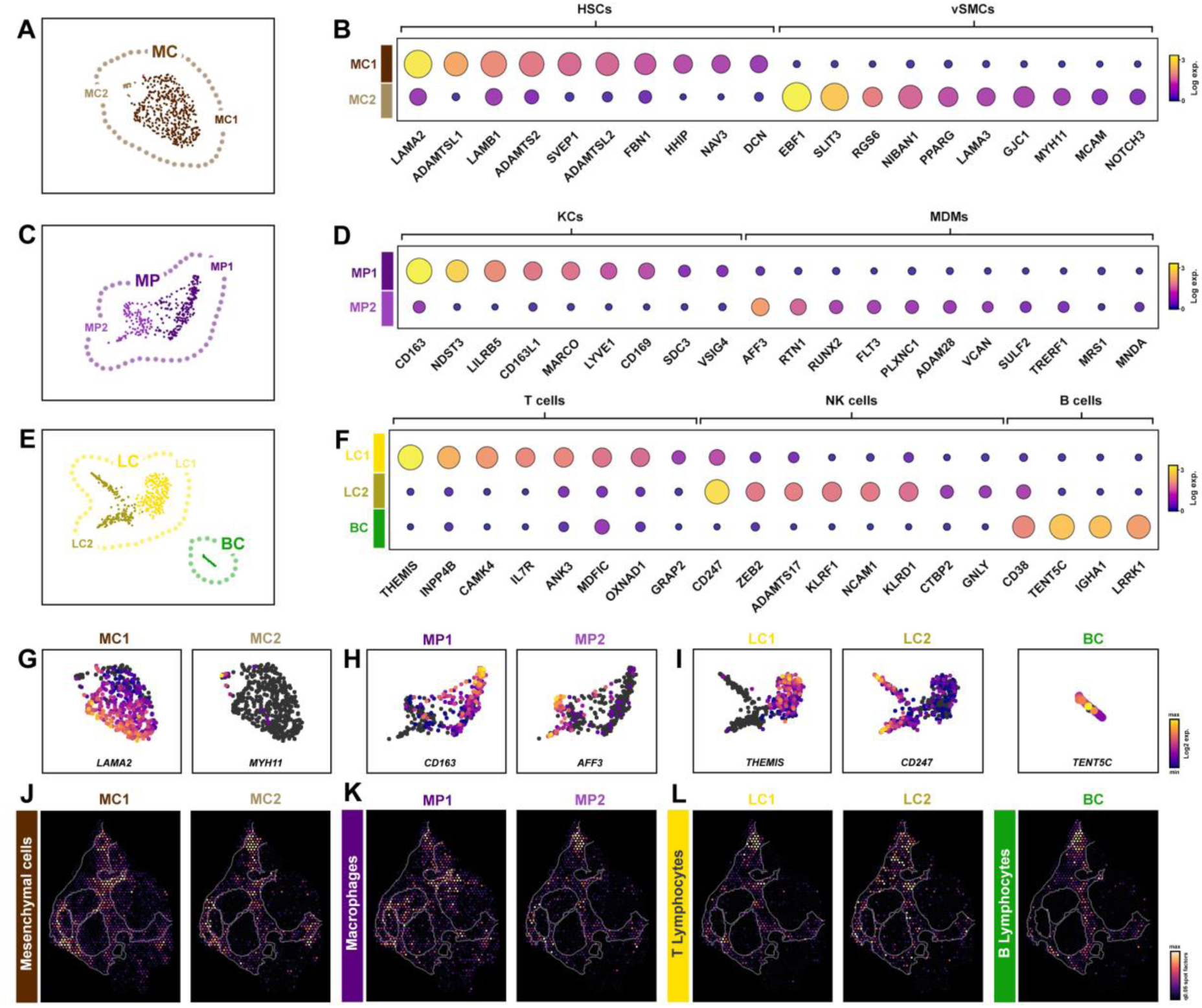
Spatial heterogeneity of non-parenchymal cell populations in cirrhotic liver. (**A**-**I**) UMAP sub-clusters, dot plots and marker gene expression plots showing heterogeneity of mesenchymal (MC; **A,B,G**), macrophage (MP; **C**,**D**,**H**) and immune (LC, BC; **E**,**F**,**I**) cell sub-populations. Spatial deconvolution maps of MC (**J**), MP (**K**) and immune (**L**) cell sub-populations within cirrhotic sample b. HSC, hepatic stellate cells; vSMCs, vascular smooth muscle cells; KCs, kupffer cells; MDMs, monocyte-derived macrophages, LC, lymphocytes; BC, b cells.

Characterisation of immune cell populations identified a broad cluster of MPs which further resolved to two distinct subpopulations (Figure 4C). MP1 was defined by *CD163*+ / *MARCO*+ cells and together with marker genes (*EXT1*, *LILRB5*, *CD169*, *NDST3*, *MSR1*, *LYVE1*) represented the liver resident macrophages, Kupffer cells (KCs) (Figure 4D, H). In contrast, MP2 was characterised by *MARCO*-/ *MNDA*+ cells and with marker genes (*AFF3*, *RTN1*, *FLT3*, *CCSER1*, *XYLT1*, *CIITA*) identified as injury-recruited monocyte derived macrophages (MDMs; Figure 4D, H)^5, 7, 9^. In keeping with these data, gene signatures of MP1 localised within the liver tissue compartment indicative of liver resident KCs, whereas MP2 was spatially distributed within the scar indicative of injury (Figure 4K). T and B lymphocyte populations resolved to three distinct clusters (LC1, LC2 and BC) (Figure 4E). Marker gene analyses detected T cell signatures in LC1 (*IL7R*, *THEMIS*, *INPP4B*, *CAMK4*, *PBX4*) while LC2 population was associated with natural killer T (NKT) cell signatures (*KLRF1*, *NCAM1*, *NCR1, LINGO2, CEP78*) (Figure 4F, I). B cells (BC) represented the smallest cluster (*IGHA1*, *TENT5C*, *LRRK1*) (Figure 4F, I). Spatial mapping of T and B lymphocytes revealed their enrichment within the scar, indicative of their role during chronic liver disease (Figure 4L). Together, these data characterised the gene signatures underlying disease-associated non-parenchymal cell populations.

### Cellular heterogeneity of parenchymal cell populations in human cirrhotic liver

To explore the gene signatures underlying parenchymal sub-clusters, we initially defined the CC population by broad cell specific marker genes (*PKHD1*, *CTNND2*, *BICC1*, *ANXA4*, *DCDC2)*. Further heterogeneity was revealed by resolving to two distinct sub-clusters (CC1, CC2; Figure 5A). Differential expression analysis showed mature CC markers (*KRT7*, *KRT19*, *CFTR, AQP1, HNF1B*) were more highly expressed in CC2, whereas interestingly CC1 demonstrated a gene signature with similarities to HCs (*GPC6*, *APOB*, *PPARGC1A*, *MYLK*, *MLXIPL*, *FGF13, FGFR3, SULF1*, *RUNX1*) (Figure 5B). Exploring these markers, *SULF1* is normally restricted to the developing fetal liver but can be upregulated in injured liver, potentially from attempted regeneration^21^. Similarly, FGF signalling is important for hepatocyte regeneration^22^ and we noted several FGF components enriched in CC1 and HCs (*FGF13*, *FGFR3*). Although the function of FGF13 in liver is less well understood, studies have indicated its potential as part of a seven-gene signature for HCC progression^23^. As a result, the profile of CC1 suggested an immature CC population that are HC-like, potentially indicative of an attempted regenerative response.

**Figure 5.**
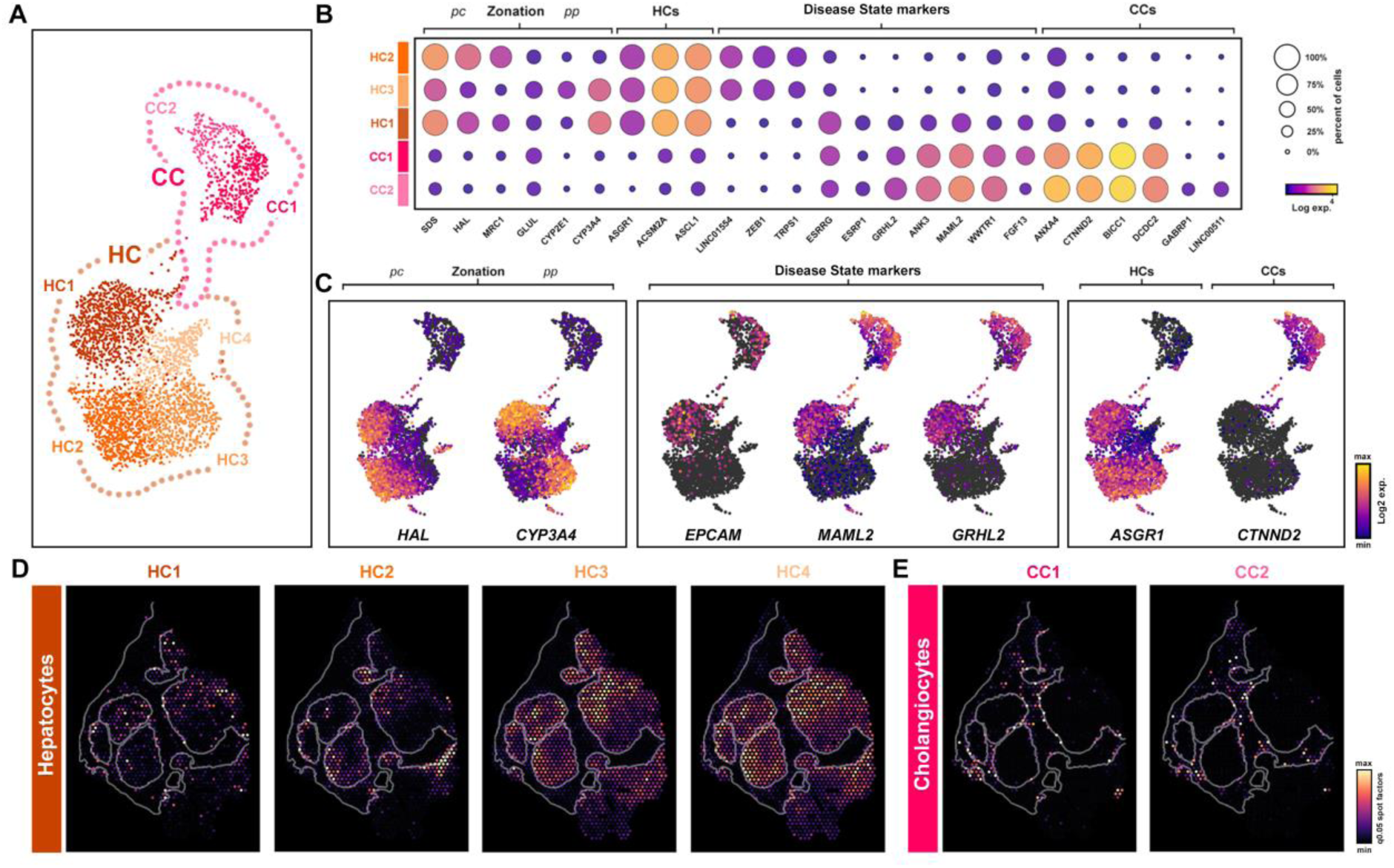
Spatial characterisation of parenchymal sub-populations in cirrhotic liver. (**A**) UMAP plot of parenchymal cells show hepatocyte (HC; HC1-HC4) and cholangiocyte (CC; CC1, CC2) sub-populations. (**B, C**) Dot and UMAP expression plots show marker genes of HC zonation, HC/CC broad cell types and disease-associated cell states. Spatial deconvolution maps of HC (**D**) and CC (**E**) sub-populations within cirrhotic sample b. PC, pericentral; PP, periportal.

HCs represented the largest cell-type cluster with 45% of sequenced nuclei. Following re-clustering, this population resolved to 4 sub-clusters (HC1-4) with HC1 isolated from HC2/3 (Figure 5A). We noted HC4 had much lower UMI counts (Supplementary Figure 5) and was not present in snATAC-seq data, therefore these cells were not characterised further. Hepatocyte zonation in the human liver has been well documented^24, 25^, including gradients of gene expression from periportal (PP) to pericentral (PC) regions which reflect distinct metabolic functions across the liver lobule. Evaluation of known PP (*HAL*, *MRC1, SDS*) and PC (*CYP3A4, CYP2E1, GLUL*) markers identified opposing gradients of gene expression with HC2/HC3 enriched for PP/PC markers respectively (Figure 5B, C). In keeping with their characteristic zonation profiles, spatial maps showed gene enrichment signatures for periportal (PP; HC2) hepatocytes were associated with portal bridging fibrosis (Figure 5D). Whereas, pericentral (PC; HC3) hepatocytes occupied complementary regions within the main tissue (Figure 5D).

HC1 displayed PP/PC zonation markers, similar to HC2/HC3, ruling out zonation as the underlying cause for segregation of this cluster (Figure 5B, C). However, further characterisation of HC1 highlighted several hepatocyte signature genes that were not present (*LINC01554*, *MAGI2*, *TRIM55*, *ZEB1*, *ZIC1*, *HEPACAM*, *TRPS1*) compared to HC2/HC3 (Figure 5B, C). Significantly, downregulation of many of these genes influences cell proliferation, migration and fate^26, 27, 28, 29, 30^ while reduced expression of *LINC01554* is associated with advancing fibrosis in NAFLD patients^31^. Further analyses revealed HC1 uniquely shared transcriptional signatures with CCs (*ARHGEF38*, *GRHL2*, *MAML2*, *ESRRG*, *ESRP1*, *ZNF83*, *EPCAM*), suggesting a transition between parenchymal cell states (Figure 5B, C). Accordingly, GRHL2 is implicated in epithelial cell fate and CC differentiation^32, 33^, while ESRP1/2 are epithelial splicing factors involved in lineage differentiation programmes and HC regeneration^34, 35, 36, 37^. Although EpCAM is not usually expressed in mature HCs, it is associated with hepatobiliary progenitors and nascent HCs in diseased liver^5, 38, 39, 40^. Significantly, *EPCAM* was uniquely expressed in HC1 identifying this cluster as potentially disease-associated, reminiscent of our previous work^40^ (Figure 5B, C). Moreover, spatially, HC1 and CC1 subpopulations were enriched at the scar-interface, while gene signatures underlying CC2 were localised further within the scar (Figure 5D, E). Taken together, snRNA-seq identified atypical HCs (HC1) that shared transcriptional signatures with immature CCs (CC1), indicating a population of disease associated parenchymal cells potentially involved in an impaired regenerative response during progressive fibrosis.

### Spatial deconvolution of the fibrotic niche

To increase our understanding of the spatial heterogeneity in cirrhotic tissue, we combined the strengths of ST and snRNA-seq gene signatures (Figure 2A and Figure 6A) using cell2location^41^ to deconvolute multi-cell ST spots (Supplementary Table 7). Using non-negative matrix factorisation (NMF), we clustered cell profiles (Figure 6B) and inferred the co-location of cell subpopulations. Spatial deconvolution of 19 cell subpopulations (Supplementary Figures 6-9) mapped to 10 co-location factors (sample b; Factor 0-9; Figure 6B), revealing several cell subpopulations spatially enriched within cirrhotic tissue (Figure 6B-D; Supplementary Figures 7 and 9). Significantly, we focussed on spatial maps that were scar associated (Factors 2, 5 & 9) and located at the scar-interface (Factors 3 & 8) (Figure 6C, D).

**Figure 6.**
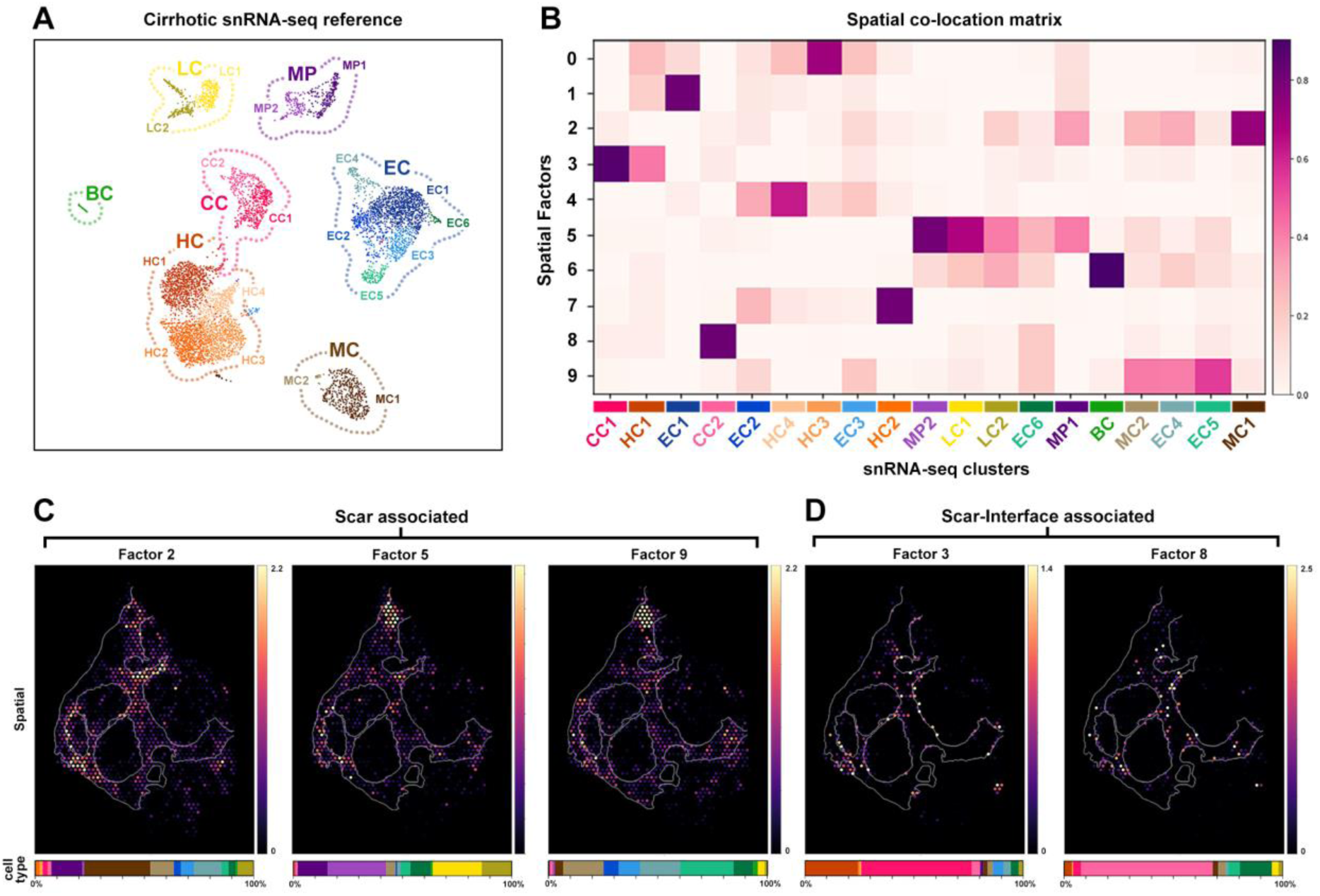
Spatial deconvolution of the fibrotic niche. (**A**) Reference gene signatures from snRNA-seq sub-populations were used to deconvolute multi-cell ST spots. (**B**) Co-location analysis for sample b using non-negative matrix factorisation (NMF) shows 10 factor maps (0-9) and the proportion of cell sub-populations to each spatial factor. (**C**) Spatial maps of scar-associated (2, 5 and 9) and scar-interface associated (**D**) factors reveal spatial co-location of cell sub-populations in cirrhotic sample b. Hepatocytes (HC), endothelial cells (EC), cholangiocytes (CC), macrophages (MP), mesenchymal cells (MC), T lymphocytes (TC), B lymphocytes (BC).

Factor 2 was comprised of HSCs (MC1), a smaller contribution from resident macrophages (MP1) and a subpopulation of ECs (EC4) (Figure 6B, C). To uncover the spatial signature of these scar-associated subpopulations, we analysed DEGs underlying Factor 2 spots and linked expression to cell-type profiles from the snRNA-seq reference (Supplementary Figure 10A, B). As expected, collagen transcripts (*COL1A1*, *COL1A2*, *COL3A1*, *COL6A2*) were significantly enriched in the scar alongside secreted ECM components and other profibrogenic markers indicative of aHSCs (MC1; *SPARC*, *AEBP1*, *TAGLN*, *FSTL3*, *LAMC3*, *RBP1*, *ADAMTSL2, IGFBP7*; Supplementary Figure 10A-C). Spatial deconvolution identified novel HSC genes, *QSOX1* and *RAMP1* (Supplementary Figure 10B, C)^17, 20^ as part of the profibrotic niche.

Several immune cell populations were spatially co-located within the fibrotic scar, represented by Factor 5 (Figure 6B, C), and comprised resident macrophages (MP1), infiltrating monocyte-derived macrophages (MDMs; MP2) and lymphocyte populations (LC1, LC2). Although these immune cells resolved to a unique factor, they co-located to similar regions of the scar occupied by aHSCs (Figure 6B, C). Spatial DEGs of Factor 5 highlighted markers of inflammatory MDMs (HLA-DQA1, HLA-DRB5, *LYZ*, *VCAN*, *CD52*, *CD74*) and T cells (IL7R, TRBC2, CD7) enriched within the fibrotic niche (Supplementary Figure 10D-F). Following injury, the pleiotropic cytokine macrophage migration inhibitory factor (MIF) is secreted and signals via cell surface receptor/co-receptors CD74/CD44 expressed on macrophages and lymphocytes^42^. Together, MIF-CD74/CD44 signalling components were spatially enriched at the scar/tissue interface, highlighting tissue regions undergoing inflammation-driven repair orchestrated by multiple immune cell-types (Supplementary Figure 10F).

Interestingly, Factor 9 demonstrated the spatial expression of vascular-associated EC subpopulations (EC4, EC5), also occupying similar hotspots within the fibrotic scar (Figure 6B, C). Spatial DEGs revealed the scar-associated expression of pericentral ECs (EC4; *PTGDS*, *TAGLN*, *ADAMTS1*, *ADAMTS4*, *TNXB*) and periportal ECs (EC5; *PLVAP*, *VIM*, *EMP1*, *VWF*, *LTBP4*) (Supplementary Figure 10G-I). In agreement with our spatial data, a recent study also verified PLVAP+ ECs as scar-associated and expanded in cirrhosis^9^.

Spatial DEGs underlying Factors 3 and 8 showed an enrichment of cholangiocyte markers (*KRT19*, *KRT7*, *CFTR, AQP1*), but also genes characteristic of liver progenitor-type cells (LPCs) (*TACSTD2*, *CD24*, *EPCAM*, *SOX9, PROM1, SPINT1, CLDN4/7, KRT23)* (Supplementary Figure 11A, B)^38^. Spatial deconvolution identified LPC markers were expressed by both hepatocyte and cholangiocyte cell subpopulations (HC1/CC1) located at the scar interface in diseased liver (Supplementary Figure 11B, D, F)^43, 44^. Significantly, KRT23 is normally restricted to CCs, however expression is upregulated in potential LPCs in cirrhotic liver^45^ and was spatially expressed at the scar interface in our cirrhotic ST data (Figure 11B, F). Similarly, the tight-junction components CLDN4 and CLDN7 are absent from HCs of healthy liver, but upregulated in hepatocytes of severely damaged human liver^46^, in agreement with our ST data (Figure 11B, F). Moreover, we have previously identified the presence of EpCAM^+^/SOX9^+^ HCs at the scar interface during progressive liver injury^40^. Although these appear reminiscent of pathways underlying the ductal plate during normal liver development, our previous data indicate increasing levels of SOX9 in these cells is a prognostic and diagnostic measure that parallels severity of liver disease. As such this is more suggestive of an impaired regenerative response due to on-going liver injury.

### Parenchymal cells show dynamic cell state transitions in cirrhotic liver

Given these data, we investigated the transcriptional dynamics of parenchymal clusters using RNA velocity and trajectory inference. Overlaying RNA velocity vectors on the snRNA-seq projection uncovered temporal dynamics within parenchymal cells and remarkably, suggested a change in cell fate from HC1 to CC1, before further differentiating to mature CCs (CC2) (Figure 7A). Using integrated snATAC-seq data, we inferred the trajectory of parenchymal cell clusters (HC3 to CC2) (Figure 7B) to highlight pseudo-temporal changes in open chromatin and gene expression associated with transitioning cell states (Figure 7C). Trajectory inference of parenchymal sub-clusters identified Notch (*MAML2*), Hippo (*WWTR1, SOX9*), FGF (*FGFR2*, *FGFR3*, *FGF13*) and WNT signalling (*BICC1*, *ESRRG*, *DCDC2, ANK3*) components were highly expressed in atypical HCs (HC1) and further increased in expression along the pseudo-time from HC1 to CC1 (Figure 7C), in agreement with our marker gene analysis (Figure 5B,C). Moreover, these data support previous studies on signalling pathways such as Wnt, Notch and Hippo in liver regeneration, cell fate decisions and differentiation^47, 48, 49, 50^, which when perturbed can lead to cirrhosis and HCC^51^. From our integrated gene regulatory data we identified a disease-associated parenchymal cell state (Figure 7C, D) expressing EPCAM, consistent with our previous work defining its localisation in an aberrant SOX9 cell population at the scar-interface in progressive liver disease^40, 52^. However, our data also highlighted *ESRP1* and *ARHGEF38* as novel genes involved in the HC1-CC1 transitional axis (Figure 7C, D).

**Figure 7.**
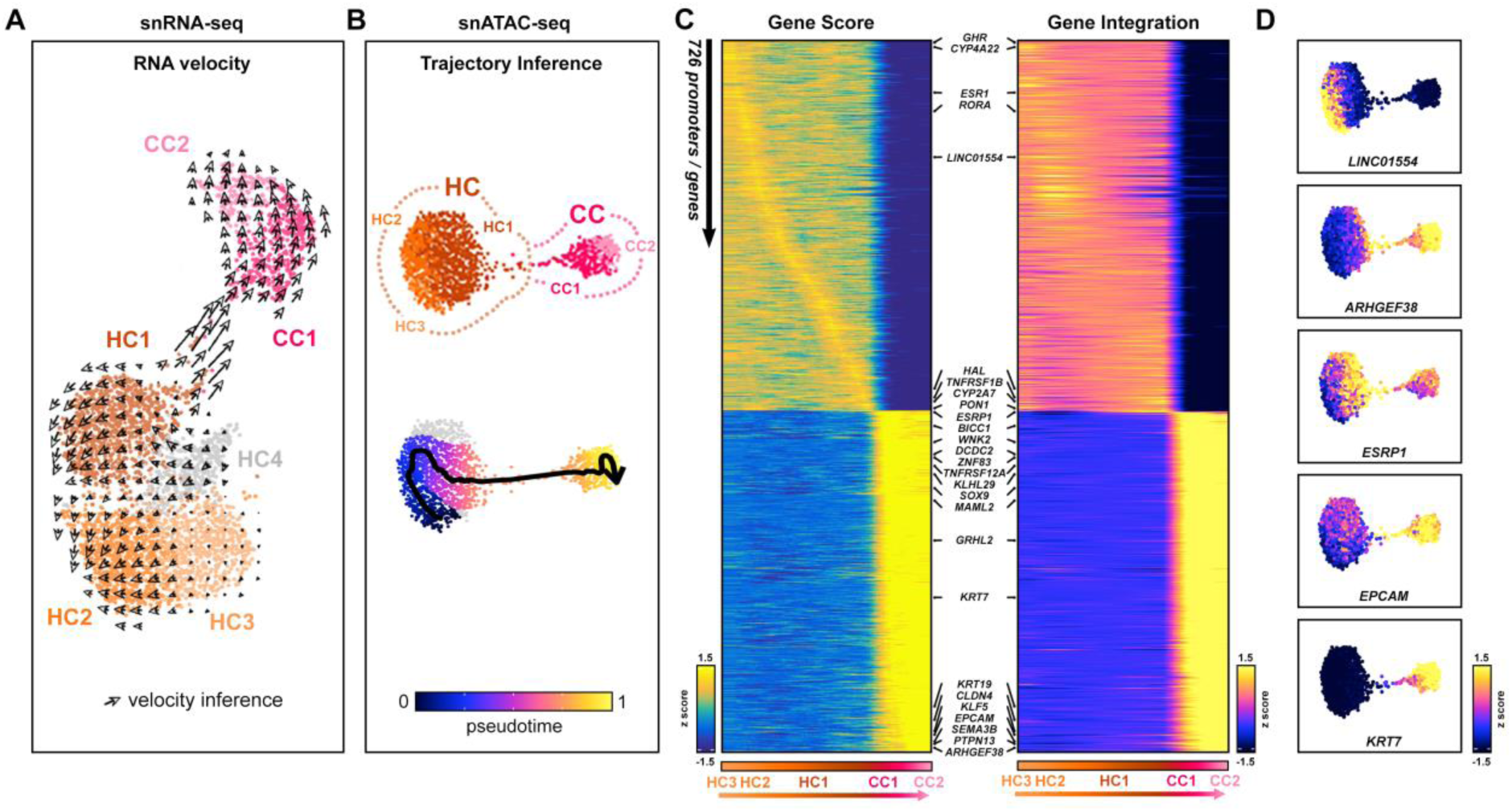
Parenchymal cells display transitional cell states and directional trajectory in cirrhotic liver. (**A**,**B**) RNA velocity (**A**) and pseudo-time (**B**) analyses infer cell state transitions from HC1 to CC1. (**C**) Integrative pseudo-time on snATAC-seq reveals correlated gene/enhancer accessibility (Gene Score) and expression (Gene Integration) of significant genes involved in the transition. (**D**) Gene Integration plots of key transition markers showing gene expression dynamics along the pseudo-time. Hepatocytes (HC), cholangiocytes (CC).

### Integrative multi-omic analysis of parenchymal cells reveal TF dynamics underlying altered cell states in cirrhotic liver

To further interrogate the gene regulatory mechanisms underlying these altered parenchymal cell states, we identified TFs whose expression correlated with changes in motif accessibility across the pseudo-time trajectory (Figure 8A and Supplementary Table 8). This approach revealed the dynamics of several core hepatic TFs (*HNF4A*, *NR5A2 (LRH-1)*, *HNF1B*, *ONECUT1*, *GATA6*, *KLF5*) associated with cell states and their potential dysregulation due to ongoing liver disease. The transition of hepatocytes (HC2/HC3) to a disease-associated state (HC1) correlated with the down-regulation of several orphan nuclear receptors (*HNF4A*, *NR1I2*/*I3*, *NR2C1*, *RORA*/*C*) (Figure 8A, B). Indeed, HNF4A is a master regulator of hepatocyte cell identity and several studies have shown its impairment in human chronic liver disease and potential as a therapeutic target in liver cirrhosis^53, 54, 55, 56, 57^.

**Figure 8.**
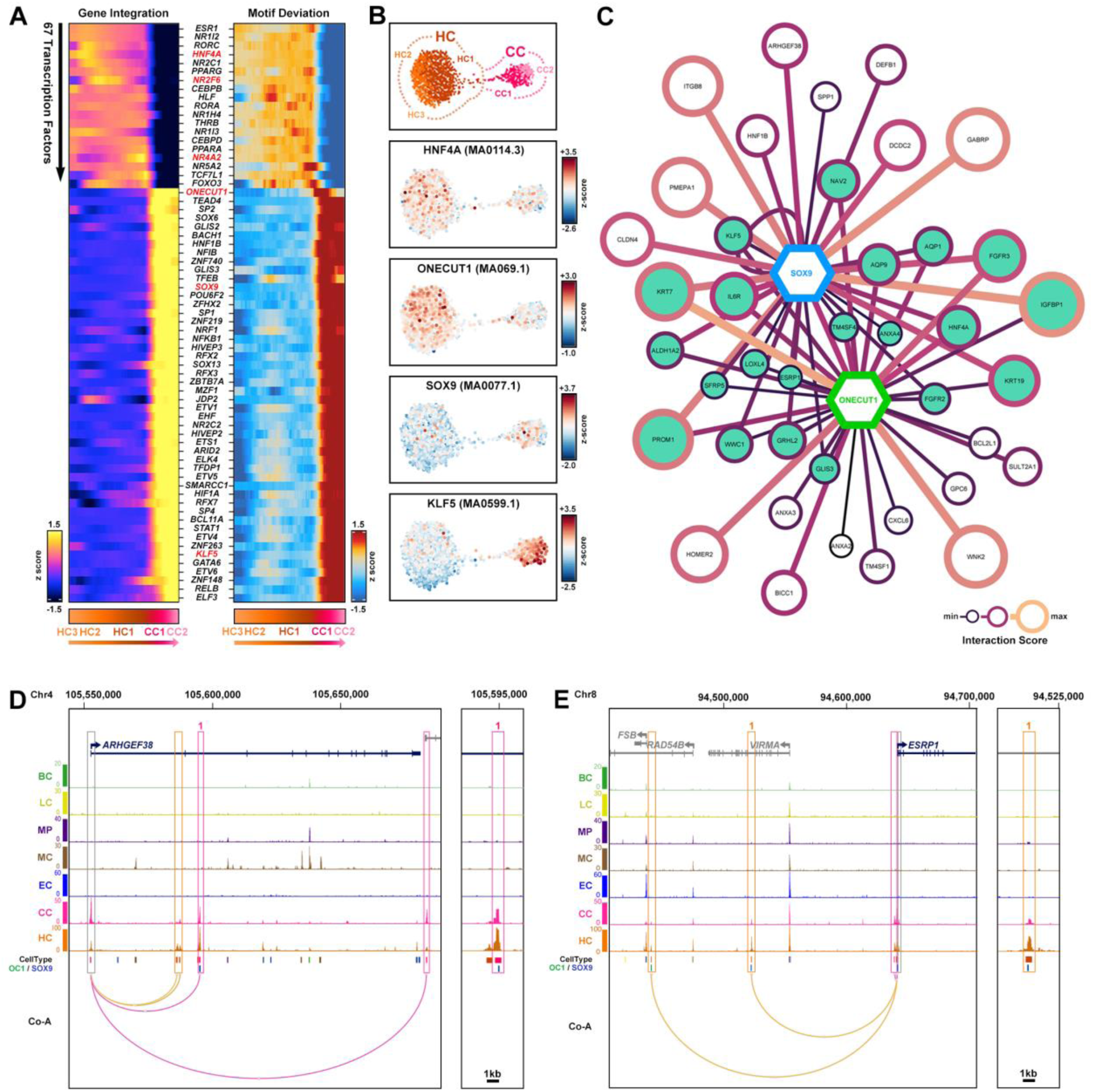
Integrative pseudo-time reveals TF dynamics underlying disease-associated parenchymal cell states at scar interface. (**A**) Integrated multi-omic analyses of parenchymal cell subpopulations (HC3-CC2) reveal the correlated expression of TFs and enhancer accessibility (motif deviation) along the transition from HCs to CCs. Select TFs of interest are highlighted (red). (**B**) UMAP motif accessibility plots (z-score) of select TFs within parenchymal cells. (**C**) Gene regulatory network (GRN) visualisation for select targets of SOX9 (blue) and ONECUT1 (green), constructed using enhancer-gene co-accessible interactions with corresponding TF motif. Potential co-regulated targets (teal). (**D**,**E**) UCSC browser screenshots of cell-type snATAC-seq tracks, show cell-type co-accessible (Co-A) enhancer-gene interactions (HC, orange; CC, pink) and TF motif locations (SOX9, blue; OC1; green) for *ARHGEF38* (**D**) and *ESRP1* (**E**). Hepatocytes (HC), endothelial cells (EC), cholangiocytes (CC), macrophages (MP), mesenchymal cells (MC), T lymphocytes (TC), B lymphocytes (BC).

Our analysis also revealed the transient expression and increased chromatin accessibility of several TFs in HC1 (*TCF7L1*, *NR5A2 (LRH-1)*, *FOXO3*) just prior to the transition to CC1 (Figure 8A). Interestingly, these TFs are implicated in Wnt signalling. TCF7L1 modulates Wnt target gene expression in the presence/absence of β-catenin, while *NR5A2* (LRH-1) itself is a β-catenin target gene with fundamental roles in fetal liver development and adult hepatocyte cell identity^56^. Conversely, FOXO3 negatively regulates β-catenin (*CTNNB1*) in hepatocellular carcinoma and supresses hepatocyte proliferation during liver regeneration^58, 59^. Collectively, our data suggests these TFs are transiently up-regulated in disease-associated cell states, and may play a role in cell fate and disease progression in cirrhosis^60^.

Consistent with our previous work, integrative TF analysis identified the downstream effector, *SOX9* across transitional cell states. Our data showed expression and enhancer accessibility in CC1, suggesting transition toward a biliary cell fate (Figure 8A, B). In addition, we also identified *ONECUT1, SOX13, RFX7* and *ZNF148* as transitional TFs. In particular we noted *ONECUT1* expression and motif accessibility spanned the transition from HC1 to CC1 (Figure 8A, B). During liver development, both SOX9 and ONECUT1 are involved in ductal plate specification. However, although SOX9 expression is maintained in mature biliary cells into adult, HNF6 is lost from the biliary lineage and expressed in SOX9 negative hepatocytes^40, 61, 62^. Given these data and gene expression profiles, we investigated how SOX9 and ONECUT1 may regulate the transitional landscape of diseased states within the HC1-CC1 axis. Consistent with their roles in ductal plate during liver development, our GRN identified SOX9 / ONECUT1 co-regulation of genes associated with a biliary and LPC phenotype (including *KRT19*, *KRT7*, *PROM1*, *FGFR2*, *TM4SF4* and *ANXA4*; Figure 8C and Supplementary Table 8)^38^. Genes involved in signalling pathways typical of cell fate transitions in hepatocytes and cholangiocytes were also highlighted (*SFRP5*, *FGFR2*, *FGFR3* and *GRHL2*) (Figure 8C)^38^. Drawing on novel genes identified in the HC1-CC1 transitional axis, our snATAC-seq data revealed co-accessible interactions for *ARHGEF38* enriched for SOX9 motifs (Figure 8C, D), whereas *ESRP1* was co-accessible with both SOX9 and ONECUT1 identified enhancers (Figure 8C, E). Although promoter activity was evident in both the HC and CC population, our snATAC-seq data contained an overabundance of transitional cell-types from HC1 and CC1 respectively (Figure 2F; Figure 8B, D, E). Moreover, peak enhancer accessibility for both *ESRP1* and *ARHGEF38* was more prevalent in the HC population, largely due to HC1 overabundance (Figure 8D, E). These data provide novel insight into liver-specific mechanisms driving disease. In particular, the transcriptional diversity driven by alternative splicing events during disease associated cell transitions is not well understood. Significantly, ESRP1 is an epithelial cell-specific RNA-binding protein that regulates alternative splicing of several genes, including *FGFR2* highlighted by our GRN and others involved in epithelial-mesenchymal transition (EMT)^63, 64^. As such, it has received much attention playing a role in tumour motility and invasiveness. Similarly, ARHGEF38, a RhoGEF implicated in tumorigenesis and metastasis^65^, is associated with cancer progression in prostate. However, these genes have more divergent roles relevant to fibrosis and potentially demarcate early transitional events in progressive disease. In particular, gene splicing is a critical aspect of gene expression driving human development and cell differentiation^64^, it seems conceivable that expression of ESRP1 in the HC1-CC1 cell populations may recapitulate fetal gene splicing to create a signature indicative of attempted regeneration. Moreover, as a direct consequence of fibrosis, mechanical properties of ECM also regulate alternative splicing, including those driven by ESRP1^66^, through effects on intracellular contractility involving RhoGEFs. In combination, this microenvironment and TF profile would conceivably alter the cell state and behaviour of our spatially identified HC1-CC1 population located at the scar interface in liver disease.

## Discussion

Single cell sequencing technologies have advanced our understanding of cellular heterogeneity in both normal and diseased human liver^5, 6, 7, 8, 9^. More recently ST studies have provided insight into the distribution of specific cell-types in healthy and diseased states, zonation in normal liver and hepatic macrophage identity in healthy and obese liver^12, 13, 14, 15^. Here we combine spatially resolved and single nucleus gene expression with chromatin accessibility to provide a comprehensive map of human liver cirrhosis and identify changing cell states associated with the scar during disease.

Our computational approaches have identified a series of spatially resolved maps to characterise changing cell states as part of the fibrotic niche in diseased liver. In keeping with their role in excessive matrix deposition, we identified specific gene signatures reminiscent of activated HSCs (aHSCs; liver specific myofibroblasts) located within the scar. Significantly these analyses also identified scar-associated expression of novel aHSC genes, *QSOX1* and *RAMP1*. QSOX1 is essential for incorporating laminin into the ECM, resulting in an aHSC response through integrin-mediated cell adhesion, migration and signalling^67, 68^. Aberrant expression of laminin and QSOX1 is also observed in various cancers^69, 70, 71^ while inhibition of QSOX1 is associated with fewer myofibroblasts, a disorganised ECM and decreased matrix stiffness in the tumour microenvironment^72^. RAMP1 is a receptor for the neuropeptide calcitonin gene-related peptide (CGRP) and functionally links sensory innervation with liver regeneration through YAP/TAZ regulation and immune infiltration^73, 74^. Thus, targeting these factors may provide a novel therapeutic route to limit progressive fibrosis and improve regenerative response.

Similar to aHSCs, several immune cell populations were also spatially resolved within the fibrotic scar, including resident macrophages, infiltrating monocyte-derived macrophages and lymphocyte populations. Significantly, the localisation and gene signatures associated with monocyte derived macrophages were indicative of an inflammatory driven response suggestive of on-going damage / repair of surrounding cells. In agreement with previous scRNA-seq studies^5, 6, 7, 8, 9^, we also spatially located a subset of scar-associated endothelial cells that were expanded and localised to regions of scarring in liver cirrhosis. These data further emphasise the multicellular response to the local environment in liver disease.

Our multi-omic approach combining spatial and single cell data provided an opportunity to investigate how hepatocyte cell states are influenced by their tissue environment. Our results highlighted a spatial pattern of gene modules specifically localised at the scar interface. Unbiased computational analysis identified these subpopulations containing genes characteristic of both hepatocytes and cholangiocytes (HC1/CC1). Previous studies aimed at understanding liver regeneration suggest that, depending on the injury, both cholangiocytes and hepatocytes are capable of functioning as facultative stem cells and transdifferentiate into either cell-type. However, the majority of these studies have been inferred from specific mouse models of severe injury^75, 76, 77, 78^. In this study, our data provided an opportunity to investigate the regenerative mechanisms and transitional states of hepatocytes and cholangiocytes directly in human. Through pseudo temporal analysis of our single nuclei data, we uncovered a directional trajectory and transitional state from hepatocytes to cholangiocytes. Mechanistically, our gene expression and transcriptional motif analysis pointed toward signalling in cirrhosis involving crosstalk between Hippo, Notch and Wnt; all perturbed and associated with liver disease^51^.

Significantly, we have previously characterised hepatocytes lining the scar in models of human and mouse liver disease and shown an impaired response with cells expressing both hepatocyte (e.g. α1AT) and cholangiocyte (CK19) markers. Moreover, as an ectopic and impaired regenerative response we have also identified these cells as EPCAM / SOX9+ (similar to ductal plate cells during human liver development)^40^. In patients with liver disease, the extent of ectopic SOX9 expressing cells in liver biopsies during early phases of liver fibrosis predicts progressive disease within 3 years^40^. As further insight into the mechanisms underpinning these EPCAM / SOX9+ altered cell states at the scar interface, we identified ONECUT1 as a critical transcriptional regulator of disease associated hepatocytes. Analysis of the integrated snRNA and ATAC-seq data identified a novel transitional axis, marked by *ESRP1* and *ARHGEF38* expression, driven by ONECUT1 and SOX9 through a co-accessibility enhancer network regulating these transitional gene states in cirrhosis. These data also highlight previously unexplored mechanisms involving ESRP1 and gene-splicing events involved in transcriptional reprogramming in progressive liver disease. These may be linked to alternative promoter use and splicing of *HNF4A* during disease, in response to both TGFβ signalling and the mechanical properties of the scar^66, 79^. Of interest, the hepatocyte gene, *HNF4A*, has two promoters and encodes at least nine isoforms through differential splicing^80^. In alcoholic hepatitis the core profibrotic cytokine TGFβ induces fetal *HNF4A*-*P2* promoter use rather than adult *HNF4A*-*P1*, resulting in dysregulated gene expression including an EMT-type response and increased levels of GRHL2^79^.

Collectively, we have provided a publicly available map and resource for further integrative studies requiring insight from single cell transcriptomics, epigenomics and spatial gene expression in human liver. Overall, these data will facilitate studies aimed at identifying new markers for early detection of liver disease, therapeutic targets to improve liver function and advance our understanding of its regenerative capacity directly during progressive disease.

## Methods

### Human tissue

Liver tissue was obtained following ethical approval and informed consent (National Research Ethics Service, REC14/NW1260/22) from the Manchester Foundation Trust Biobank. Fresh, unfixed tissue was obtained intraoperatively from patients undergoing surgical liver resections at the Manchester Royal Infirmary. Tissue was collected from three independent patients. All had early cirrhosis (Childs-Pugh A), confirmed histologically by an expert clinical histopathologist in the non-malignant regions of the resection. Patients were as follows: sample a, 71 year old male undergoing hepatectomy for large HCC; sample b, 68 year old male undergoing liver resection for hepatic metastatic colon cancer; sample c, 58 year old male undergoing resection for HCC. Tissue was snap frozen within 30 minutes of collection and stored at -80°C.

### Sample processing, library preparation and sequencing

Snap frozen liver tissue was rapidly thawed in a 37°C water bath for ∼60 seconds. Tissue was minced on ice with sterile razor blades alongside addition of cold lysis buffer (10mM Tris HCl, 10mM NaCl, 3mM MgCl_2_, 0.1% IGEPAL, 0.1% Tween20, 0.01% Digitonin, 0.2U/ul RNAse inhibitor). Minced tissue was transferred to a falcon tube and incubated on ice in lysis buffer for 1h with periodic gentle vortexing every 10-15 minutes. Tissue was passed through 40um and 20um filters and centrifuged at 500g for 10 minutes at 4°C. The pellet was gently resuspended in wash buffer (10mM Tris HCl, 10mM NaCl, 3mM MgCl_2_, 0.1% Tween20, 0.2U/ul RNAse inhibitor). The suspension was split in half to provide nuclei for independent snATAC-seq and snRNA-seq runs. The centrifugation and wash step was repeated before a final centrifugation at 500g for 5 minutes at 4°C. The pellets were then resuspended in either 0.1% ultrapure BSA in PBS plus 0.2U/ul RNAse inhibitor for snRNA-seq, or 1x nuclei buffer diluted from 20x nuclei buffer (10x Genomics) for snATAC-seq.

Nuclei were processed through Chromium Next GEM Single Cell ATAC and Chromium Next GEM Single Cell 3’ protocols by the University of Manchester Genomic Technologies Core Facility. Briefly, for snATAC-seq nuclei suspensions were incubated in a Transposition Mix to fragment accessible DNA and ligate adapter sequences to the fragments prior to GEM generation and barcoding, cleanup and library construction. For snRNA-seq, GEMs were generated for barcoding and reverse transcription. cDNA was amplified and a library generated after fragmentation, end repair and A-tailing and adapter ligation steps. Sequencing was carried out on the Illumina NextSeq 500 Platform.

### snRNA-seq data processing and clustering

#### Data pre-processing

The sequence files generated from the sequencer were processed using 10x Genomics custom pipeline Cell Ranger v3.1.0^81^. The fastq files generated from the pipeline are then aligned to the hg38 custom genome with all the default parameters of Cell Ranger. Cell Ranger identifies the valid barcodes with nuclei and counted UMIs associated to each nucleus. It uses STAR aligner to align reads to genome while discarding all the counts mapping to multiple loci during counting. The uniquely mapped UMI counts are reported in a gene by cell count matrix represented as sparse matrix format. Cell Ranger outputs 9,754 nuclei which is then taken for further downstream QC.

#### Filtering

The low-quality cells were removed from the dataset, to ensure that the technical noise does not affect the downstream analysis. We looked into three commonly used parameters for cell quality evaluation, the number of UMIs per cell barcode (library size), the number of genes per cell barcode and the proportion of UMIs that are mapped to mitochondrial genes^82^. Cells that have lower UMI counts than 0.5 Median Absolute Deviation (MAD) for these three parameters were filtered out. To check whether there are cells that have outlier distributions, which can indicate doublets or multiplets of cells, violin plots were used on these three parameters. The outlier cells that have total read counts more than 100,000 were removed, as potential doublets. After filtering, 6170 nuclei remained for downstream analysis.

#### Classification of cell-cycle phase

The cyclone method was used to calculate for each cell the score of G1 phase and G2M phase^83^. Cyclone uses a pre-build model that is trained on a known cell-cycle dataset where the sign of difference in the expression between a pair of genes was used to mark the cell-cycle stages. The cell-cycle phase of each cell is identified based on the observed sign for each marker pair of each of the phases.

#### Normalization

To make sure that the counts are comparable among cells, we applied library-size normalization, where each cell is normalized by the total counts over all the genes in the dataset. The total counts were set to 10,000 reads per cell. Logarithmic transformation (natural log) was then applied to the data matrix.

#### Visualization & Clustering

As a first step for visualization and clustering the highly variable genes (HVG) were identified using Scanpy’s *highly_variable_genes()* function^84^. This function identifies HVGs by implementing a dispersion-based method. In this approach the normalized dispersion is calculated by first grouping the genes in multiple different bins and then scaling with the mean and standard deviation of the gene dispersion for which the mean expression falls into a given bin. These HVG genes were then used to reduce the dimensions of the dataset using PCA. The dimension of the dataset was further reduced to 2D using t-SNE and UMAP with 40 PCA components. To calculate the UMAP, a neighborhood graph was first generated of cells using Scanpy’s *neighbors()* function with *the n_neighbors* parameter set to 10. Then, the UMAP was generated using this neighborhood graph. For clustering, the SNN (Shared Neighborhood Network) method was used, which identified 7 clusters in the dataset.

#### Identification of marker genes

To identify marker genes for each cluster, Scanpy’s *rank_gene_groups()* function with the Wilcoxon method was used. Differentially expressed genes are reported based on a comparison of each cluster’s marker genes against all other clusters. These marker genes were then used to annotate the cell types of a cluster.

#### RNA-velocity

scvelo v0.2.4 was used to calculate the RNA-velocity of the data^85^. The spliced vs un-spliced ratio was calculated using the velocyto CLI method. This method gives a ratio of 18% spliced vs 82% un-spliced for this data. The default stochastic method to calculate the velocity was used.

### snATAC-seq data processing and clustering

#### Data pre-processing

snATAC-seq fastq files were processed using 10X Genomics custom pipeline for snATAC-seq, cellranger-atac v.1.2.0 and was mapped to hg38 custom genome with all the default parameters. The pipeline identified 4,743 nuclei with 6,238 median fragments per nucleus. With about 42.5% of fragments overlapped called peaks.

#### Cell Quality Control

To conduct quality control of snATAC-seq data, two parameters were considered, the number of unique nuclear fragments and the signal-to-backround ratio and the signal-to-background ratio. The former is represented as the fragments that are not mapping to mitochondrial DNA, while the latter, is measured as TSS enrichment score, which is calculated as the ratio between the peak enrichment at transcription start site raltive to the flaning region. Cells having lower than 1000 ATAC-seq fragments or lower than 4 TSS enrichment are filtered out. 4,680 cells passed this filtering for downstream analysis.

Potential doublets were identified using ArchR’s *addDoubletScores()* function^86^. This function first synthesizes in-silico doublets and then cells that are within these synthetic doublets’s neighborhood in the UMAP embedding are identified as potential doublets. This procedure is iterated for thousands of times to calculate the confidence on a cell being a potential doublet. To identify these doublets, the default parameters were used, where 219 of 4680 cells (4.7%) were identified as potential doublets and were filtered out.

#### Dimensionality Reduction and Clustering

The iterative Latent Semantic Indexing (LSI) was applied on a genome wide 500-bp sized tiles^86^. Initially the iterative LSI identifies lower resolution clusters that mostly correspond to major cell-types. Then, the average accessiblity of each peak across these clusters is computed and being used in order to identify peaks that are most variable across the dataset. This information is used in the subsequent iteration for the identification of the informative clusters. To cluster the cells, ArchR’s *addClusters()* function was used, which is based on a graph clustering implemented in Seurat^18^, where clusters are identified in LSI sub-space. To visualize our snATAC-seq data, a 2D UMAP embedding generated with ArchR’s *addUMAP()* function was used.

#### Cell annotation and peak calling

Cells from snATAC-seq were annotated by integration with our snRNA-seq data. Cells from snATAC-seq were aligned to cells from snRNA-seq by comparing gene expression data from snRNA-seq with the gene score data from snATAC-seq. The gene score was calculated by counting the accessiblity within the gene body and 100 KB upstream/downstream of the Transcription Start Site (TSS), weighted based on the distance from the TSS. Cell type labels from the aligned snRNA-seq are then transferred to the corresponding snATAC-seq cells. The gene expression from the aligned snRNA-seq are also assigned to snATAC-seq data as a pseudo-gene expression value.

As the snATAC-seq data are essentially binary, peaks can not be called on individual single-cells. Therefore, pseudo-bulk replicates were defined based on clusters and called peaks on these pseudo-bulk data. Macs2 was utilised for peak calling and, for the downstream analysis, a 501 bp fixed width peak was used^87^. To identify marker peaks for each cluster, the *getMarkerFeatures()* function was used, which compares each cell group to its background. These marker peaks are then used to conduct motif enrichment anlaylsis for each cell-type. For motif annotations, CIS-BP (Catalog of Inferred Sequence Binding Preferences) database was used^87^. Peaks that are linked to genes, were identified by using the *addPeak2GeneLinks()* function. To infer these links, this function looks at the correlation between the peak accessibility and the gene expression, which gives the potential regulatory regions for each peak.

#### Trajectory Analysis

To order cells in a pseudo-time, ArchR aligns cells in N-dimensional LSI subspace based on a user-defined trajectory backbone. The mean cordinates for each cluster in N-dimensional suspace are then calculated and only the cells for which the euclidean distance between these mean cordinates are within the top 5% of all cells are retained. The cells are then ordered based on the distance from the cell’s own cluster and its next cluster mention in the backbone. These cells are then aligned along the embedding. Finally ArchR scales the alignment to 100. The changes in GeneScore and GeneIntegration matrix across pseudo-time were visualized using heatmap plots based on *plotTrajectoryHeatmap()* function. To have integrated pseudotime heatmaps, the *correlateTrajectories()* function was used, to identify features that changes in a correlated manner in two different matrices.

### Gene regulatory network inference

Using snATAC-seq data, all significant (correlation >0.45) co-accessible peak2gene interactions were identified and enhancers were annotated to cell-type clusters. From this library, we extracted all parenchymal specific (HC/CC) interactions associated with genes in the HC-CC pseudotime transition and scanned enhancer peaks for SOX9 (MA0077.1) and ONECUT1 (MA0679.1) motifs using FIMO^88^. Utilizing this catalog of potential SOX9 and/or ONECUT1 regulated targets, we constructed GRNs using Cytoscape v3.9.1^89^ for up to 40 genes of interest, visualizing node size/colour and edge thickness/colour based on interaction significance score (-LOG10(FDR)).

### Visium ST sample preparation, optimisation, library preparation and sequencing

Snap frozen cirrhotic liver resections were embedded in optimal cutting temperature (OCT), frozen and cryosectioned with 10μm thickness at -10°C (Leica CM1900) with a chamber temperature of -20°C. Sections were transferred onto capture areas of a chilled Visium Tissue Optimisation Slide and stored for 24hrs at -80°C before continuing with the manufacturer protocol the following day. Tissue sections were permeabilized, captured mRNA was reverse transcribed using fluorescently labelled oligonucleotides and imaged *in situ* using a high resolution slide scanner with TRITC filters (3D-Histech Pannoramic-250). A tissue permeabilization timeseries determined 12 minutes was the optimum permeabilization duration for human liver tissue (Supplementary Figure 2).

Visium Spatial Gene Expression slides were prepared according to the manufacturer protocol. Tissue was fixed, permeabilized (12 mins), mRNA captured, stained for hematoxylin and eosin (H&E) and imaged for brightfield on a high-resolution slide scanner (3D-Histech Pannoramic-250). Spatial libraries were constructed, QC’d, and sequenced at depth on NextSeq500 platform (Illumina). Following sequencing, spatial reads were assigned back to the corresponding tissue section using Space Ranger software and the data explored using Loupe software and other bioinformatic tools. All samples were sequenced to a mean depth of 255.3M reads, detecting on average 2,986 unique genes and 11,995 UMIs per spot, with the entire dataset comprising a total of 8,028 spots across four tissue sections, including one technical replicate (Supplementary Figure 3).

### Visium ST data processing and clustering

Space Ranger analysis pipelines were used to process Visium Gene Expression data. For each sample, the fiducial frames from H&E images were manually aligned within Space Ranger, spots over tissue selected and processed using the standard fresh-frozen sample workflow. UMI counts were handled by Space Ranger, prefiltering the data to remove duplicates and sequencing errors in UMIs. Regions with unusually low UMIs due to sectioning/processing artefacts were reviewed after the workflow and if deemed necessary, the sample re-processed omitting these spots.

Dimensionality reduction was handled by Space Ranger, using Principle Component Analysis (PCA). The data was visualised in 2D by passing PCA-reduced data into t-Stochastic Neighbor Embedding (t-SNE)^90^ and Uniform Manifold Approximation and Projection (UMAP)^91^ algorithms, both plotted and visualised within Loupe.

Space Ranger performed Graph-based (Louvain)^92^ and K-means clustering (across a range of values) to group spots by expression similarity. For each clustering method, differentially expressed genes (DEGs) per cluster were identified relative to the rest of the sample using the Loupe browser. Gene ontology (GO) analysis was performed using Enrichr^93^ on the top 50 DEGs per cluster. GO data was presented as a manhatten-style dot plot and significant GO terms were selected per cluster. GO terms were mapped spatially by plotting the average expression signature (Log2 average UMIs) of 5 significantly enriched genes per term. Spatial gene expression was visualised within Loupe browser. For each gene, violin plots were used to set upper and lower UMI thresholds before visualising.

### Spatial cell-type deconvolution and colocation analysis

To map the spatial distribution of individual cell-types and infer cell-type proportions for each spatial spot, we used cell2location^41^; a python tool based on Bayesian modelling. This tool was selected in preference to other methods due to its high accuracy and computational efficiency. Cell2location estimates relative and absolute cell abundance at each spatial spot by decomposing the gene expression counts in ST data into a set of reference signatures (cell-types). Reference signatures were derived prior to deconvolution analysis by computing the average expression of each gene in each cell cluster. To perform deconvolution analysis, ST data, snRNA-seq data and cell-type subpopulation clusters were used as inputs to cell2location. Genes with zero number of counts were removed from the snRNA-seq data and further filtered such that only genes expressed by at least 3% of cells were retained. Furthermore, for genes with high counts in non-zero cells, only those expressed by at least 0.05% of cells were retained. As recommended, this ensured rare cell-type populations were retained but only when expressed at high levels. Cell2location was trained using the following parameters and priors: number of training iterations: 20000; prior on the expected number of cells per location (N): 8; prior on the expected number of cell-types per location (A): 5; prior on the expected number of co-located cell-type groups per location (Y): 2; prior on the sensitivity of spatial technology: μ = ½, σ² = ⅛. Cell2location was also used for spatial co-location analysis of cell subpopulations. This was performed via a non-negative matrix factorisation (NMF) method using the estimated cell abundances as input.

To identify DEGs for each factor, we determined an appropriate minimum expression threshold for spots per factor (herein referred to as hotspots), imported spots above threshold into Loupe browser and computed DEGs versus all spots. Significant spatial genes were subsequently annotated to snRNA-seq expression data, showing the proportion of reads per gene across cell subpopulation clusters as a heatmap.

## Contributions

KPH conceived and designed experiments. NLH, SMB, SG, AA, MJD, NB, ADS, NAH, MR contributed to experimental design. VSA, AKS, HVMS provided reagents. NLH, AA, EJ, KS, VSA performed experiments. NLH, SMB, SG, AA, EJ, MR and KPH analyzed the data. NLH, SMB, SG performed computational analysis. SMB, SG, VSA, MJD, NB, ADS, NAH and MR contributed to editing and writing the manuscript. KPH and NLH guided experiments, analysed data and wrote the manuscript.

## Acknowledgements

This work was supported by the Medical Research Council (MRC; KPH, MR/J003352/1 & MR/P023541/1; NAH, MR/000638/1 & MR/S036121/1). The Genomics Core Facility and the Bioimaging Facility at the University of Manchester are acknowledged for their technical help and support from the Wellcome Trust (105610). This work was supported by the Wellcome Trust Institutional Strategic Support Fund (204796). KPH is a member of the Wellcome Trust supported Centre for Cell-Matrix Research (203128/Z/16/Z) and receives funding from the Centre’s Helen Muir Fund.

## Competing financial interests

The authors declare no competing financial interests.

## Notes

### Competing Interest Statement

The authors have declared no competing interest.

