## Supplemental Information for "A spatial map of human liver cirrhosis reveals the patho-architecture and gene signatures associated with cell state transitions during liver disease"

**Profiling the spatial transcriptional landscape defines tissue architecture in human liver cirrhosis**

To investigate the spatial transcriptional landscape of human liver cirrhosis fresh, unfixed tissue from three clinically cirrhotic donors were cryosectioned and mounted onto capture areas of a spatially barcoded Visium ST slide (Supplementary Figures 1A and 2). Tissue sections were stained for hematoxylin and eosin (H&E) and imaged at high resolution before library preparation and sequencing (Supplementary Figure 1A). Following processing and mapping of the raw sequencing data, an average of 2,986 genes were detected and mapped to 11,995 unique molecular identifiers (UMIs) per spot on the ST arrays. The entire dataset comprised a total of 8,028 spots across four tissue sections (including one technical replicate; Supplementary Figure 3). Following manual annotation, histological landmarks were apparent demarcating fibrous scar from the surrounding liver parenchyma. As expected for human samples the degree of fibrosis varied, ranging from numerous fibrous septa (sample a) to extensive bridging septa encompassing regenerative nodules (sample b) and localised portal fibrosis (sample c) (Supplementary Figure 1B-D).

To explore gene expression underlying the cirrhotic samples, dimensionality reduction of spatial data was performed followed by unsupervised clustering and differential gene analysis (Supplementary Figure 1A). Initially, we used *k*-means clustering to investigate spatial gene expression. Projecting spots and their clusters back onto the samples revealed spatial spots that could resolve fibrotic landmarks from the surrounding tissue (Supplementary Figure 1B-G). Localised regions of fibrosis were present in sample c and this was reflected by a single cluster (c2; orange) (Supplementary Figure 1D, G). However, in samples with more extensive scarring (samples a, b), several clusters were associated with fibrotic scars (Supplementary Figure 1B, C). Here, spatial clustering suggested heterogeneity within the scar, such that tissue at the scar interface was defined by clusters a2, b2 (orange) while the centre of the scar was mainly represented by clusters a4, b4 (red) (Supplementary Figure 1E, F). Interestingly, cluster b5 (purple) defined discrete locations towards the edge of the scar and was only present in the sample containing more extensive fibrosis (Supplementary Figure 1F). In contrast, *k*-means clusters broadly represented the functioning liver parenchyma by 1-2 large clusters per sample (a1, a3; b1; c1, c2) (Supplementary Figure 1B-D).

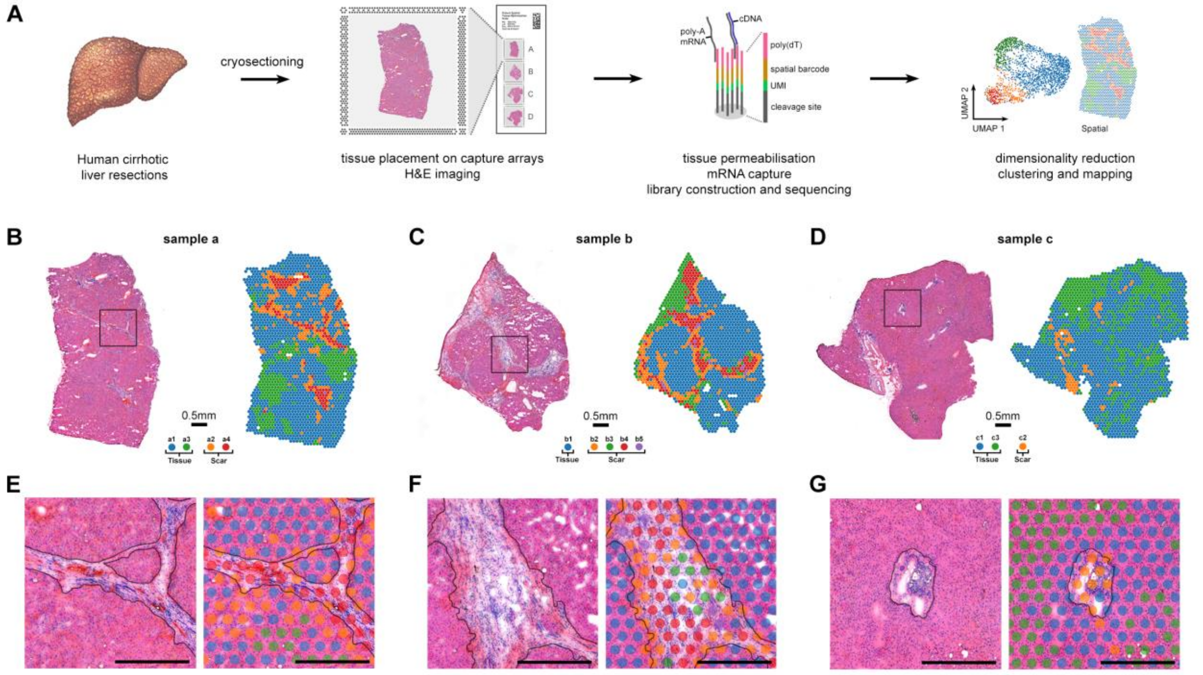


**Supplementary Figure 1. ST resolves the patho-architecture of human cirrhotic liver.** (**A**) Schematic of the Visium spatial transcriptomics (ST) workflow. (**B-D**) H&E stained cryosections and the associated spatial cluster assignments (*k*-means) for three human cirrhotic liver samples. (**E-G**) High resolution projections of clustered spots from indicated regions (box) define liver parenchyma (blue) from fibrotic scars (red) and the interface between them (orange). Scale bar, 500 µm.

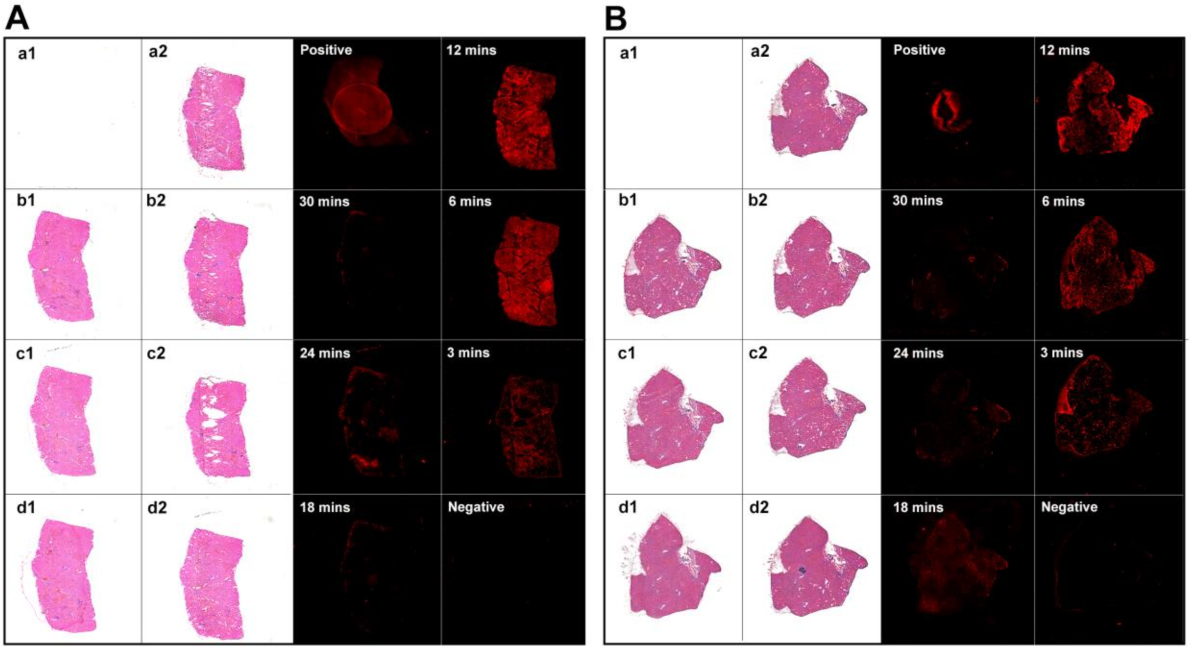


**Supplementary Figure 2. Visium Tissue Optimisation Assay.** (**A**, **B**) Serial cryosections from two independent human liver samples were placed on capture areas of Visium Optimisation slides (**A**; a1-d2, **B**; a1-d2) and imaged following histological staining. A permeabilization timeseries (0-30 mins) followed by fluorescent cDNA synthesis and imaging, shows the peak fluorescent signal is achieved after 12 minutes. Positive control, mouse cDNA; negative control, minus permeabilization enzyme.

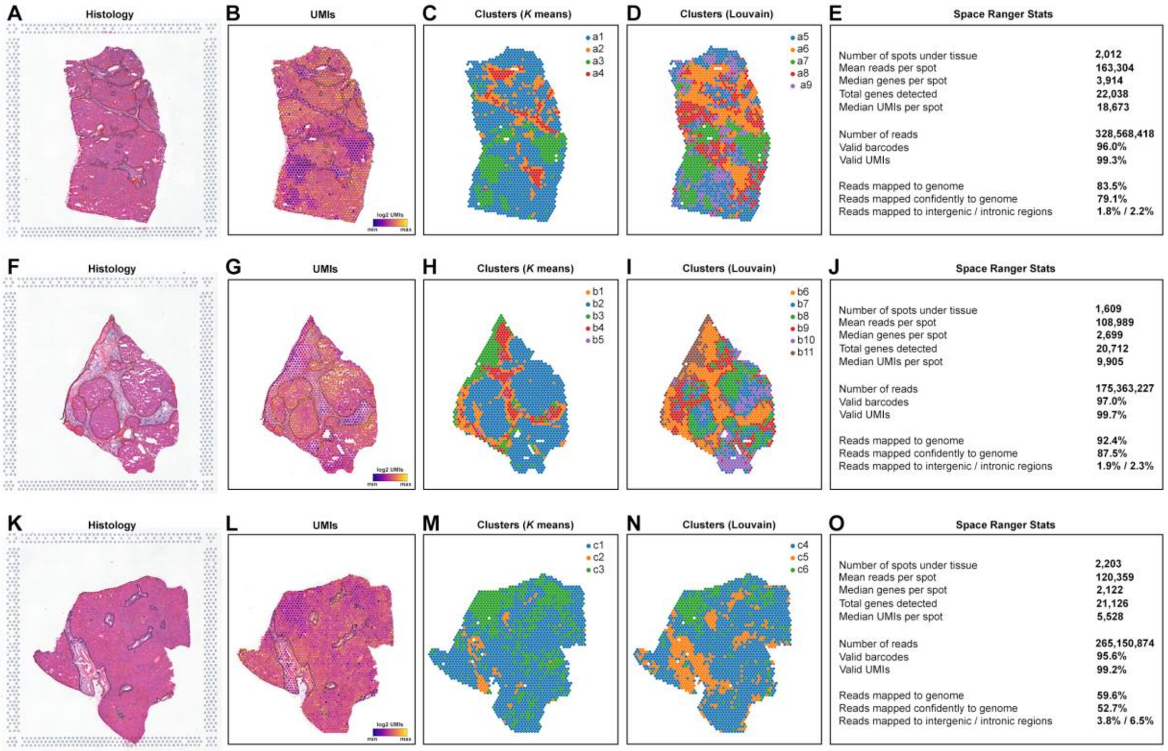


**Supplementary Figure 3. Visium Spatial transcriptomics data for human cirrhotic liver.** (**A**-**O**) Summary of spatial mapping data for cirrhotic liver *sample a* (**A**-**E**), *sample b* (**F**-**J**) and *sample c* (**K**-**O**). Brightfield imaging (**A**, **F**, **K**), spatial mapping of unique molecular identifiers (UMIs) (**B**, **G**, **H**), spatial clusters (**C**, **D**, **H**, **I**, **M**, **N**) and Space Ranger mapping statistics (**E**, **J**, **O**) for each sample.

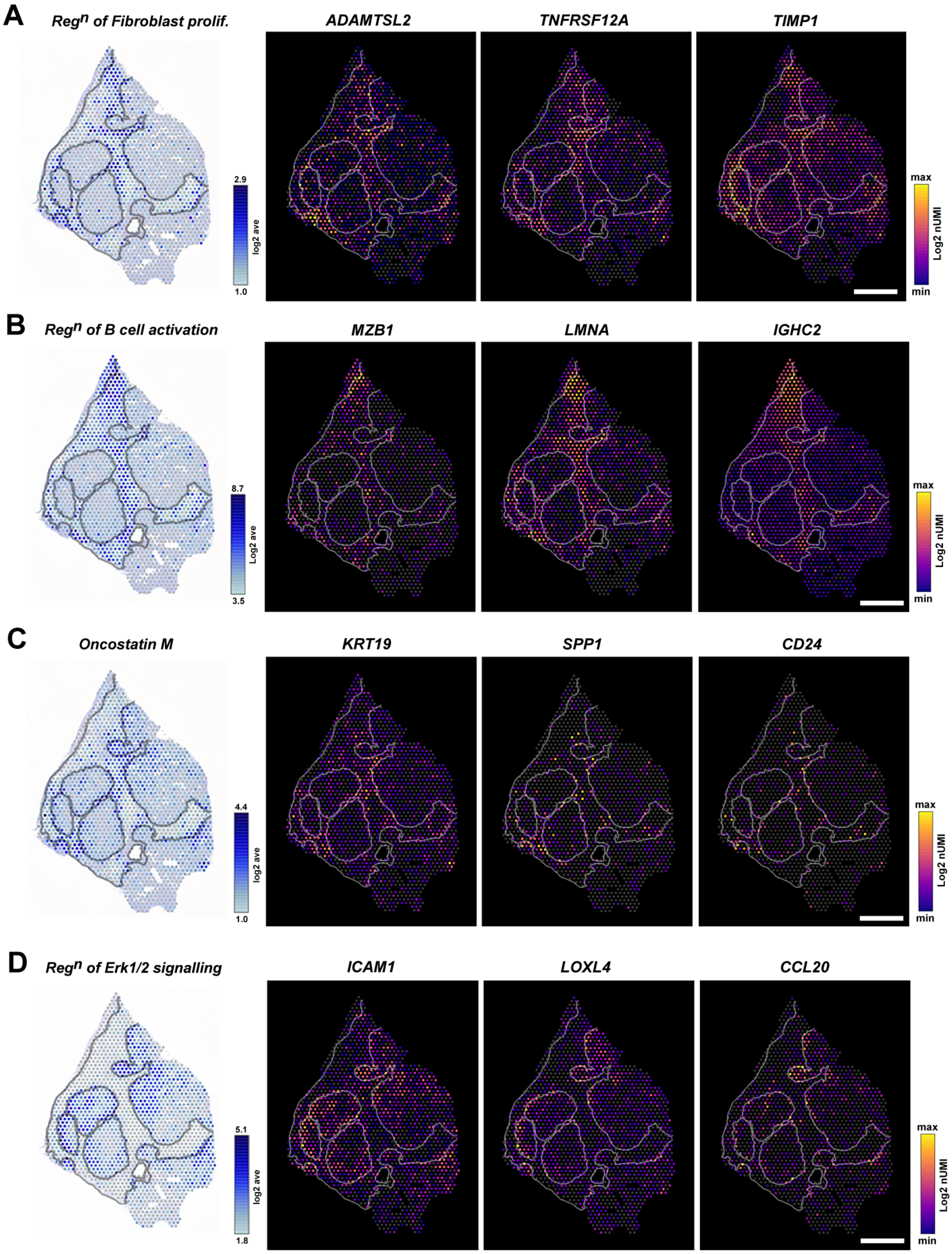


**Supplementary Figure 4. Spatial mapping of scar associated gene ontology terms .** (**A-D**) Spatial expression of gene modules underlying GO terms with corresponding spatial expression of select genes. Scale bars, 1mm.

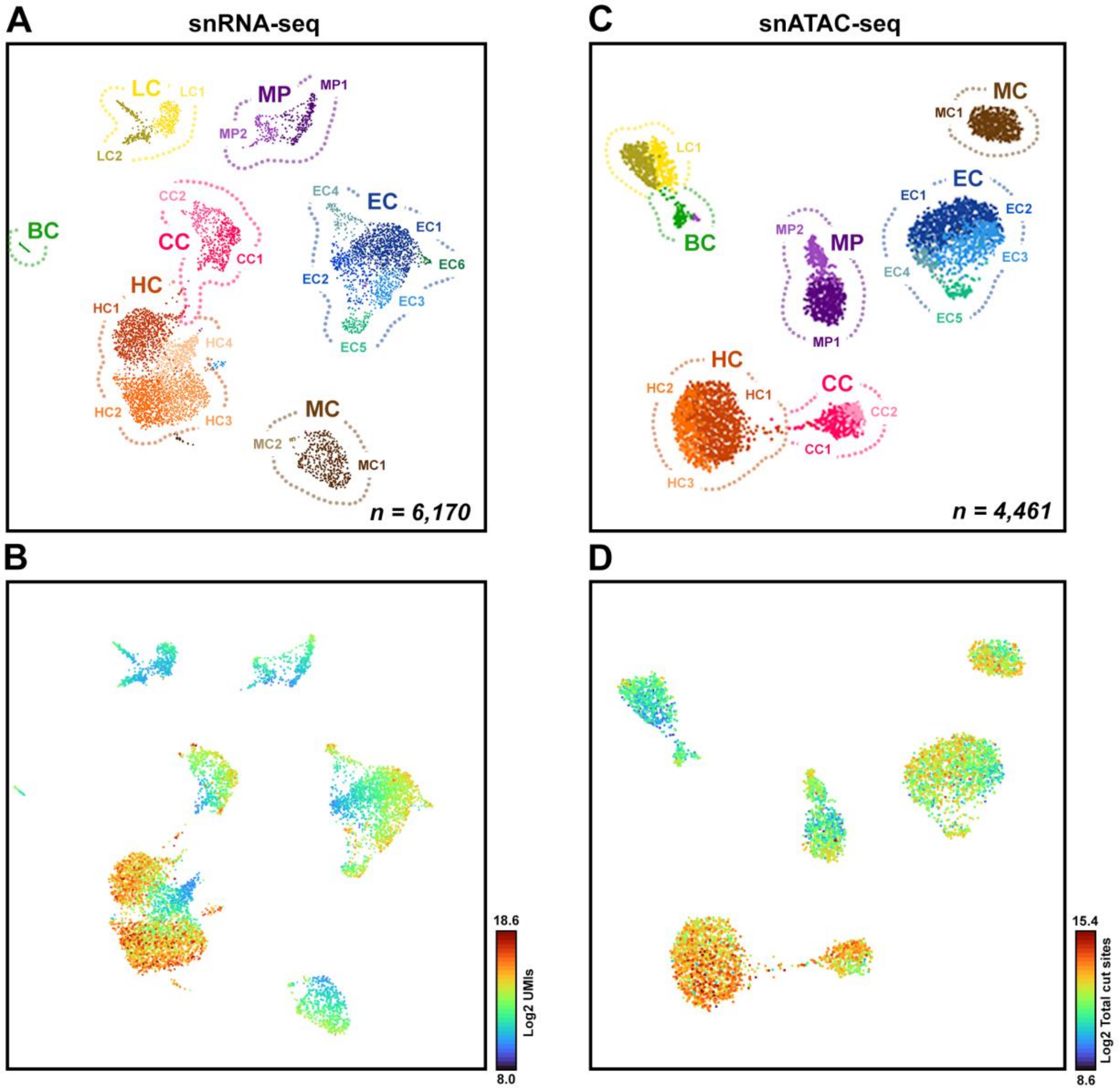


**Supplementary Figure 5. snRNA-seq and snATAC-seq count metrics** (**A**,**B**) UMAP plots of snRNA-seq sub-clusters and associated unique molecular identifier counts per cell (Log2 UMIs). (**C**,**D**) UMAP plots of snATAC-seq sub-clusters and associated total cut sites per cell. Hepatocytes (HC), endothelial cells (EC), cholangiocytes (CC), macrophages (MP), mesenchymal cells (MC), T lymphocytes (TC), B lymphocytes (BC)

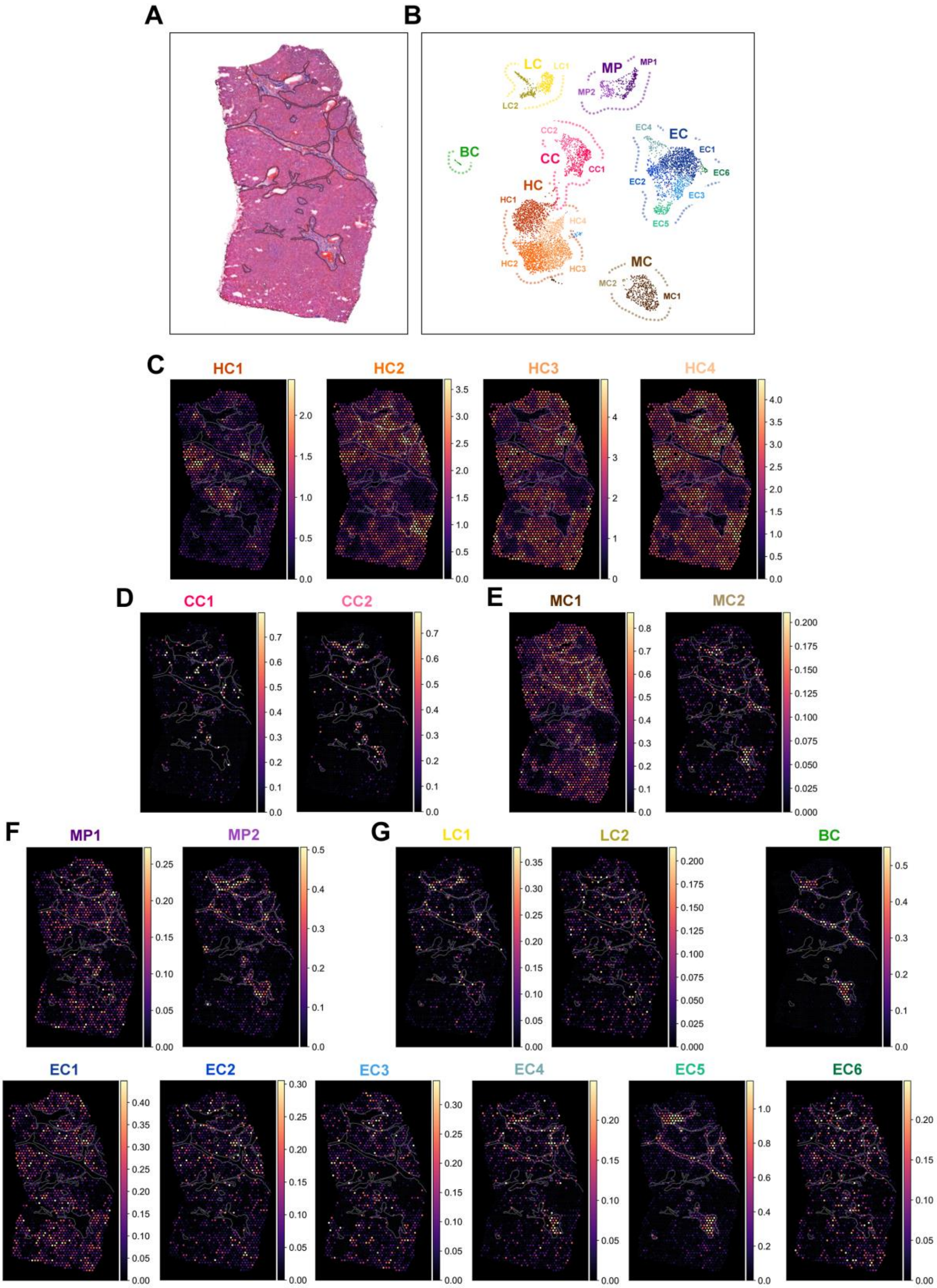


**Supplementary Figure 6. Deconvolution and spatial mapping of cell type sub-clusters in sample A.** (**A**) Brightfield imaging of sample a, annotated to show fibrotic scars. (**B**) snRNA-seq cirrhotic liver cell type sub-clusters were spatially mapped using cell2location, showing regions occupied by hepatocytes (**C**; HC1-4), cholangiocytes (**D**; CC1-2; **D**), mesenchymal cells (**E**; MC1-2), macrophages (**F**; MP1-2), T and B lymphocytes (**G**; LC1-2, BC) and endothelial cells (**H**; EC1-6).

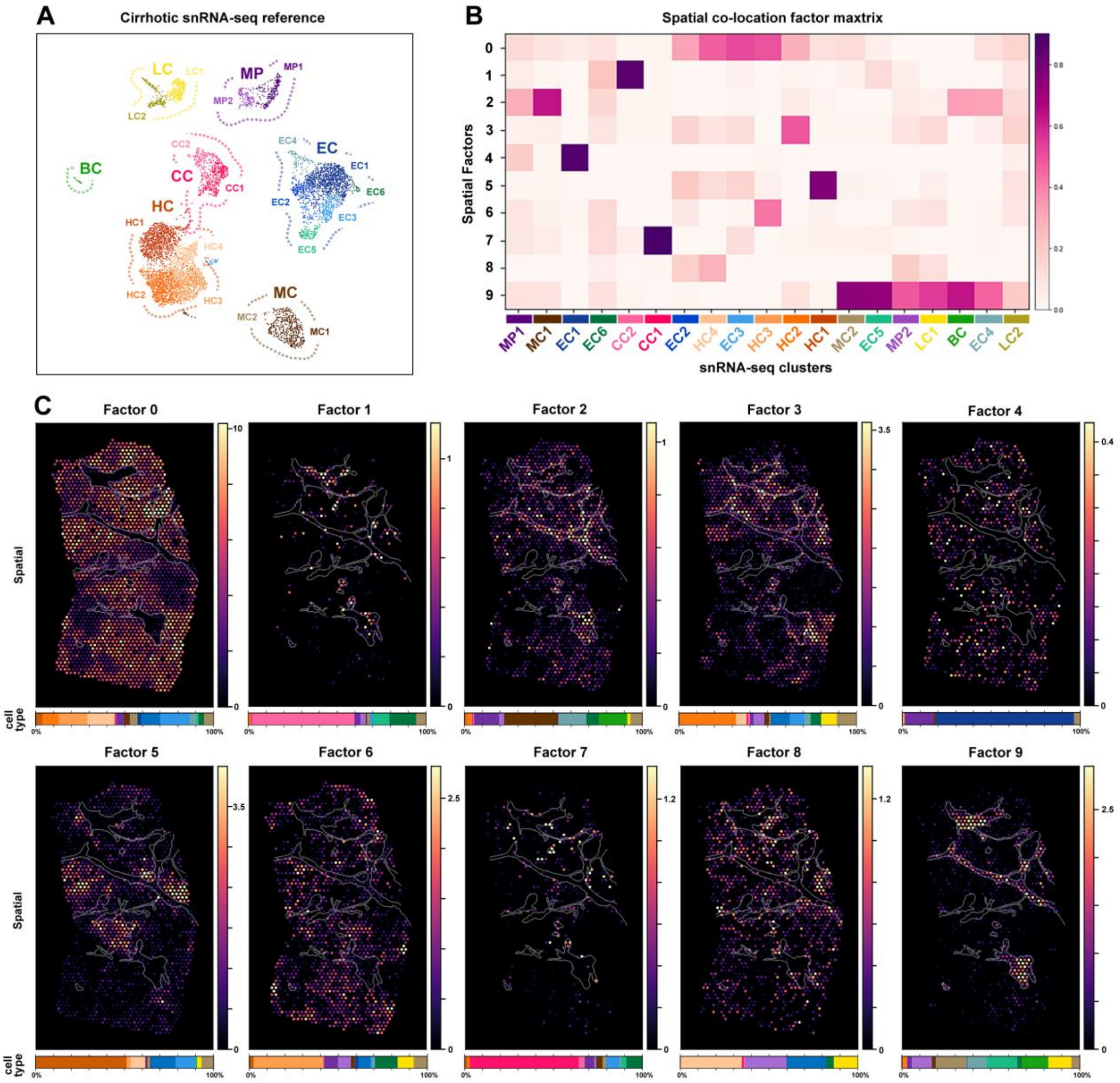


**Supplementary Figure 7. Co-location analysis of cell type sub-clusters in sample A.** (**A**) Reference gene signatures from snRNA-seq subpopulations were used to deconvolute multi-cell ST spots. (**B**) Co-location analysis using non-negative matrix factorisation (NMF) shows 10 factor maps (0-9) and the proportion of cell sub-clusters represented by each spatial factor. (**C**) Spatial mapping of co-location factors (0-9) with bar charts showing percentage of cell subpopulations which represent each factor. Hepatocytes (HC), endothelial cells (EC), cholangiocytes (CC), macrophages (MP), mesenchymal cells (MC), T lymphocytes (TC), B lymphocytes (BC).

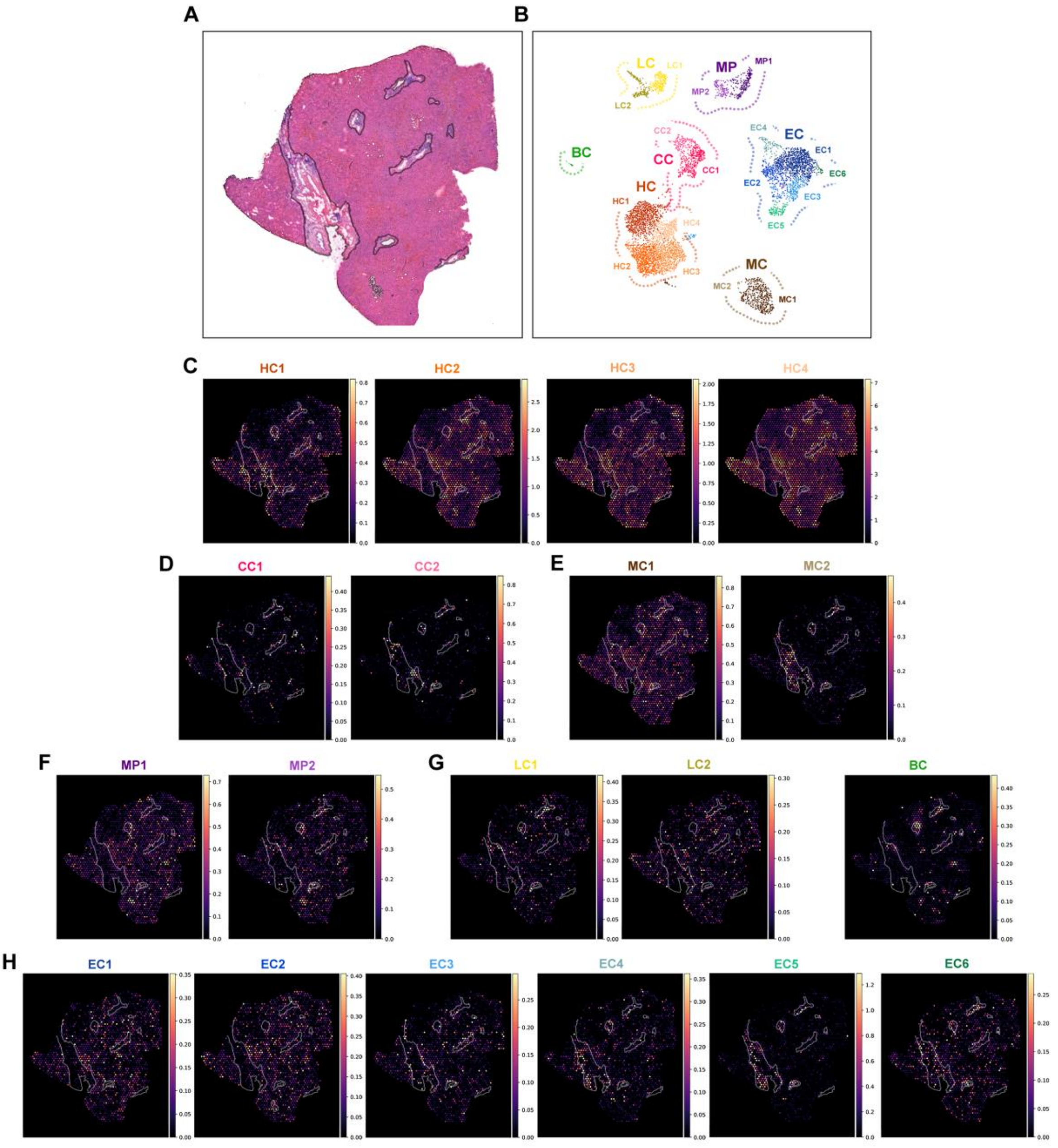


**Supplementary Figure 8. Deconvolution and spatial mapping of cell type sub-clusters in sample C.** (**A**) Brightfield imaging of sample c, annotated to show fibrotic scars. (**B**) snRNA-seq cirrhotic liver cell type sub-clusters were spatially mapped using cell2location, showing regions occupied by hepatocytes (**C**; HC1-4), cholangiocytes (**D**; CC1-2; **D**), mesenchymal cells (**E**; MC1-2), macrophages (**F**; MP1-2), T and B lymphocytes (**G**; LC1-2, BC) and endothelial cells (**H**; EC1-6).

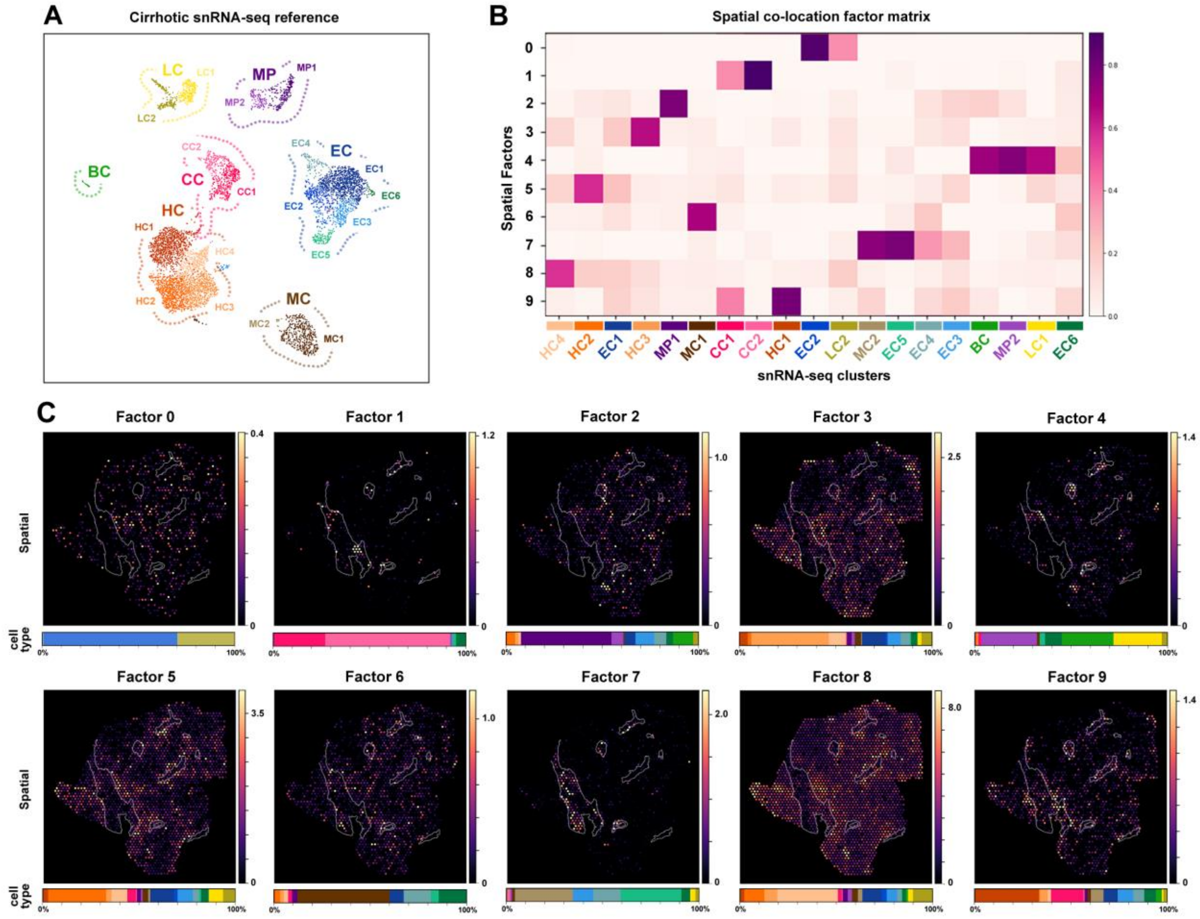


**Supplementary Figure 9. Co-location analysis of cell type sub-clusters in sample C.** (**A**) Reference gene signatures from snRNA-seq subpopulations were used to deconvolute multi-cell ST spots. (**B**) Co-location analysis using non-negative matrix factorisation (NMF) shows 10 factor maps (0-9) and the proportion of cell sub-clusters represented by each spatial factor. (**C**) Spatial mapping of co-location factors (0-9) with bar charts showing percentage of cell subpopulations which represent each factor. Hepatocytes (HC), endothelial cells (EC), cholangiocytes (CC), macrophages (MP), mesenchymal cells (MC), T lymphocytes (TC), B lymphocytes (BC).

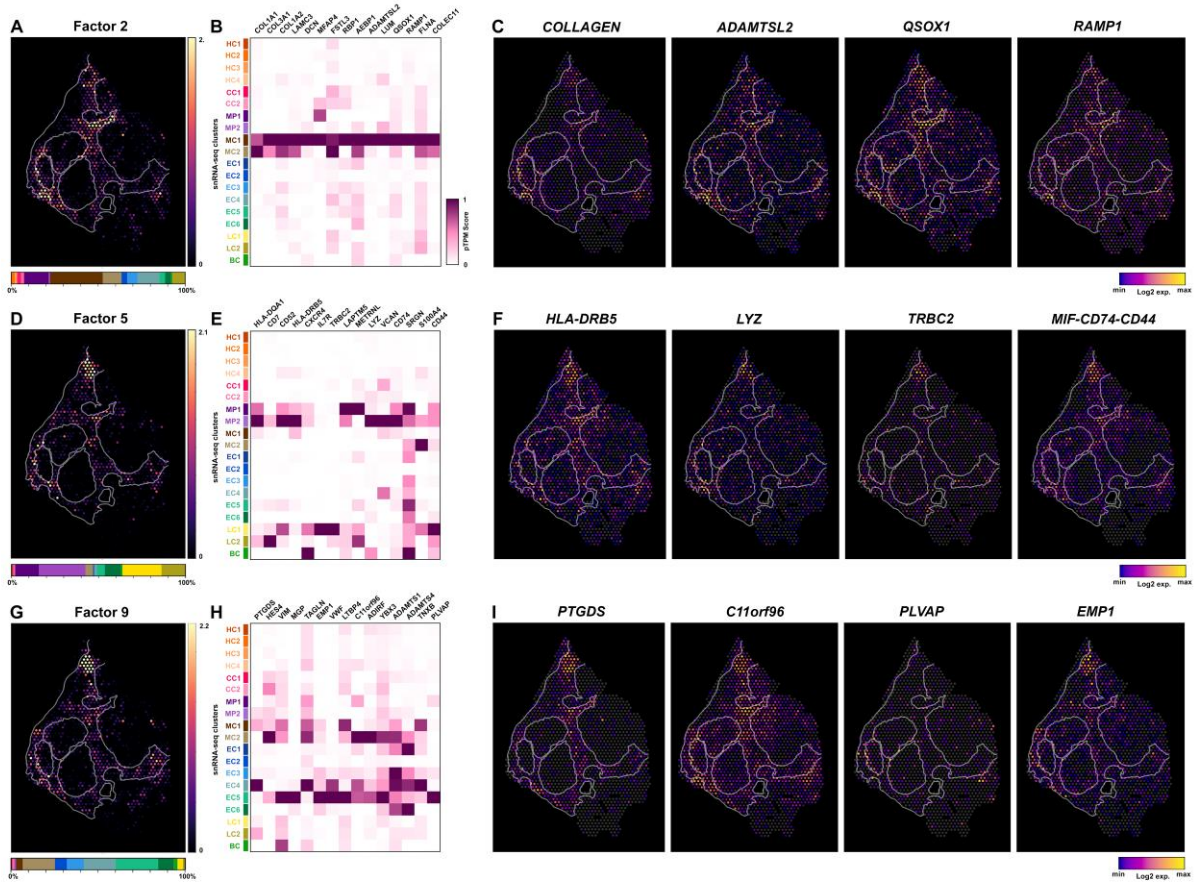


**Supplementary Figure 10. Non-parenchymal cell sub-populations are co-localised within the fibrotic niche.** (**A-I**) Spatial deconvolution of non-parenchymal cell types show several sub-populations are co-localised (Factors 2, 5 and 9) within the fibrotic scar (**A**,**D**,**G**). Selected DEGs of Factors 2, 5 and 9 (**B**,**E**,**H**) are correlated with their snRNA-seq expression score and reveal the spatial expression of scar-associated targets (**C**,**F**,**I**). Hepatocytes (HC), endothelial cells (EC), cholangiocytes (CC), macrophages (MP), mesenchymal cells (MC), T lymphocytes (TC), B lymphocytes (BC).

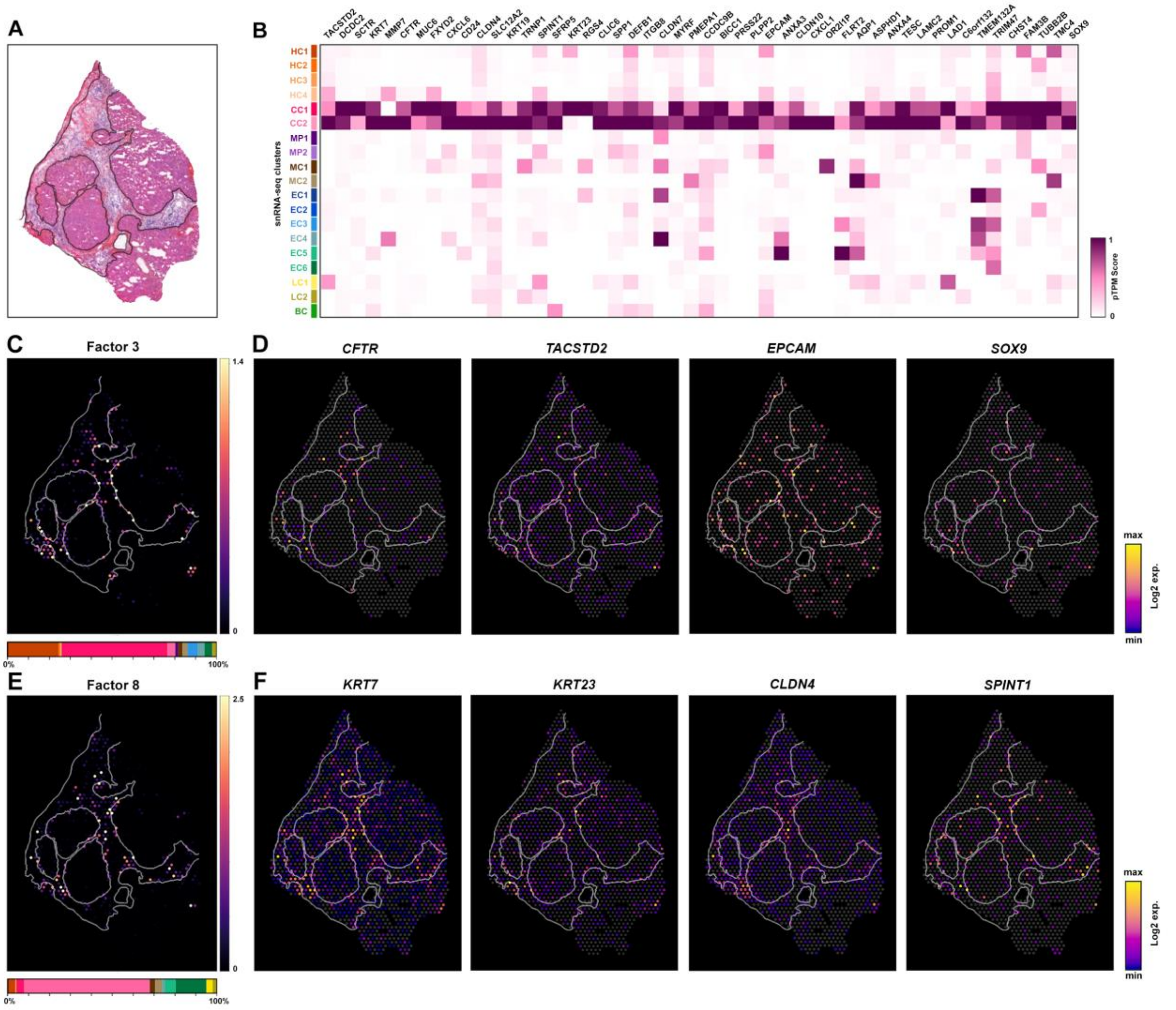


**Supplementary Figure 11. Disease-associated hybrid parenchymal cells are co-localised to the scar-interface**. (**A-F**) Spatial deconvolution of parenchymal sub-populations reveal HC1/CC1 cells are co-localised at the scar-interface (**C**; Factor 3) whereas CC2 sub-populations are preferentially located within and at the scar edge (**E**; Factor 8). Selected DEGs underlying Factors 3 and 8 (**B**) reveal spatial expression of liver progenitor cell (LPC) and cholangiocyte markers enriched at scar-interface (**D**,**F**). Hepatocytes (HC), endothelial cells (EC), cholangiocytes (CC), macrophages (MP), mesenchymal cells (MC), T lymphocytes (TC), B lymphocytes (BC).
